# Pervasive RNA Binding Protein Enrichment on TAD Boundaries Regulates TAD Organization

**DOI:** 10.1101/2023.11.22.567635

**Authors:** Qiang Sun, Qin Zhou, Yulong Qiao, Hao Sun, Huating Wang

**Affiliations:** Department of Orthopaedics and Traumatology, Li Ka Shing Institute of Health Sciences, The Chinese University of Hong Kong, Hong Kong, China; Center for Neuromusculoskeletal Restorative Medicine, Hong Kong Science Park, New Territories, Hong Kong SAR, China; Department of Chemical Pathology, Li Ka Shing Institute of Health Sciences, The Chinese University of Hong Kong, Hong Kong, China

**Keywords:** RBP, 3D genome, TAD, transcription, RBFox2

## Abstract

Mammalian genome is hierarchically organized by CTCF and cohesin through loop extrusion mechanism to facilitate the organization of topologically associating domains (TADs). Mounting evidence suggests additional factors/mechanisms exist to orchestrate TAD formation and maintenance. In this study, we investigate the potential role of RNA binding proteins (RBPs) in TAD organization. By integrated analyses of global RBP binding and 3D genome mapping profiles from both K562 and HepG2 cells, our study unveils the prevalent enrichment of RBPs on TAD boundaries and define boundary associated RBPs (baRBPs). We also characterize chromatin features of baRBP binding and uncover clustering among baRBPs and with transcription factors (TFs). Moreover, we found that baRBP binding is correlated with enhanced TAD insulation strength and in a CTCF independent manner. Further analyses revealed that baRBP binding is associated with nascent promoter transcription thus RBP/transcription may synergistically demarcate TADs. Additional experimental testing was performed using RBFox2 as a paradigm. Knockdown of *RBFox2* in K562 cells causes remarkable TAD reorganization and boundary loss. Moreover, we found RBFox2 enrichment on TAD boundaries is a conserved phenomenon in C2C12 myoblast (MB) cells. RBFox2 is down-regulated and its bound boundaries are remodeled during MB differentiation into myotubes (MTs). Knockout of *Rbfox2* in MBs also causes significant boundary reorganization. Finally, transcriptional inhibition in C2C12 cells indeed decreases RBFox2 binding and disrupts TAD boundary insulation. Altogether, our findings demonstrate that RBPs can play active role in modulating TAD organization through co-transcriptional association and synergistic action with nascent promoter transcripts.

## Introduction

The genome is packed into the tiny nuclear space in three-dimensional (3D) arrangement through proper chromatin contacts at multi-scale levels [1]. It is segregated into two compartments at multi-megabase (Mb) scale, termed A and B compartments. At a finer scale, the chromatin is organized into self-interacting domains termed the topologically associating domains (TADs), which range between 0.2-1.0 Mb. At an even finer scale of several to hundreds of kilobase (Kb) within the TADs, sub-TADs and chromatin loops can be observed [2–5]. TADs exert transcriptional influence by insulating promoter-enhancer interactions thus TAD integrity is critical for proper gene expression. Loss of a TAD boundary can result in gene mis-expression because of inappropriate enhancer-promoter (E-P) interactions. Deletion or inversion of boundary elements often alters the insulation strength of self-interacting TADs and may enable crosstalk between two neighboring TADs, which often results in aberrant gene regulation [6–9]. The role of TADs in genome regulation has evolved into a major inquiry within the field of genome biology in the last decade. Despite its privileged role in gene regulation, the mechanisms underpinning the formation and maintenance of TADs remain obscure [10–12]. It is traditionally thought to be driven by architectural proteins in contact with DNA that can anchor DNA-DNA interactions. In particular, the main force behind TAD organization is thought to be loop extrusion, a mechanism involving protein complexes comprising members of the structural maintenance of chromosome (SMC) protein family, such as cohesin, that thread double-stranded DNA (dsDNA) strands [13]. The end result is DNA loops with DNA-DNA interactions occurring at the bases of these loops [14, 15]. Cohesin extrusion is further blocked, with cohesin complexes stabilized, at TAD boundaries by the presence of the CCCTC-binding factor (CTCF), an architectural protein. Despite its essential role in vertebrates, CTCF does not seem to be required for TAD formation in some species, indicating CTCF-independent mechanisms may also generate TAD domains [16]. Interestingly, the boundaries of these domains are commonly featured by the presence of actively transcribed RNAs. Burgeoning evidence supports that active promoters or chromatin features linked to them may demarcate TADs. For example, in mammals, a subset of TAD boundaries overlaps with actively transcribed genes but not with CTCF binding sites [6]. Also, in *Drosophila. melanogaster*, most of TAD boundaries overlap with active promoters rather than with CTCF sites [17]. It is further speculated that the transcription machinery or an open transcription bubble can participate in TAD formation. However, transcription perturbation studies employing transcription inhibition in *Drosophila* embryos had little effect on TAD formation [18]. Also, forced activation of promoters in mouse embryonic stem cells (mESCs) by tethering a strong transcription activator was not enough to recapitulate the formation of a TAD boundary [7]. Therefore, it remains a puzzle whether and how an actively transcribed gene can behave as a boundary in a similar manner to a CTCF element. Recent two years have witnessed a fervent search for the answer. For example, Calandrelli R. et.al. showed that chromatin-associated RNAs (caRNAs), which include the nascent transcripts and fully transcribed RNAs that are retained on or recruited to chromatin, promote chromatin looping in multiple cells. It triggers an interesting idea that caRNAs may trap chromatin binding proteins or RNA binding proteins (RBPs) to facilitate boundary formation.

After transcription, RNA transcripts are immediately bound by RBPs that modulate all aspects of RNA metabolism and processing at post-transcriptional levels [20–22]. Many RNA-processing events are thus tightly coupled with transcription, highlighting the functional integration of transcriptional and post-transcriptional machineries. It is foreseeable that RBPs may participate in such integration processes thus triggering the fervent in elucidating RBP functions beyond its traditional framework. Indeed, increasing evidence suggests that many RBPs have direct roles in transcriptional regulation. Notably, to broadly investigate the potential function of RBPs at chromatin levels, Xiao et. al. [23] recently conducted a systematic ChIP-seq survey of a myriad of RBPs in human hepatoblastoma cell line HepG2 and chronic myelogenous leukemia cell line K562; this breakthrough work reveals the prevalence of RBP association with chromatins and a general preference for gene promoters. Findings from our own recent study [24] also shows a classical RBP, hnRNPL, interacts with enhancer transcribed nascent transcripts, eRNAs, to modulate promoter activation. Furthermore, a recent study from Wen et. al. shows that in mESCs, RBPs constitute almost one third of the chromatin associated proteins [25], among these so called chromatin-bound RBPs (chrRBPs).

In relation to 3D genome organization, a few exciting reports demonstrate the potential partaking of RBPs in 3D chromatin topological organization. For example, disruption of *Hnrnpu*, an mRNA splicing regulator, causes compartment switch and decreases loop intensity in mouse hepatocytes [26]. AGO1, which is classically known to bind with small RNAs, is found to be enriched at TAD boundaries; depletion of *AGO1* causes compartment switch, increased number and decreased size of TADs [27]; AGO1 also binds at active enhancer regions which is eRNA dependent. It is recently demonstrated that 3’end RNA processing factors, including nuclear exosome and Pfs2, co-localize with cohesin and facilitate the 3D genome organization in both yeast and mESCs [28]. These individual based studies highlight the previously unaware participation of these RBPs in 3D genome organization; However, many questions remain to be answered, for example, what are the underlying mechanisms? How extensively RBPs participate in the process?

To fill the gap, in this study, we harnessed the rich datasets of RBP ChIP-seq and Hi-C from K562 and HepG2 cells and performed integrated analyses to uncover that a wealth number of RBPs are enriched at TAD boundaries and thus defined boundary associated RBPs (baRBPs). We found the enrichment is associated with increased insulation strength and in a CTCF independent manner. Further analyses demonstrate that baRBP binding at TAD boundaries is associated with nascent promoter transcription. Using RBFox2 as a paradigm, we performed in-depth experiments to show that depletion of *RBFox2* indeed disrupts TAD formation and inhibition of transcription attenuates RBFox2 binding at chromatin and impairs TAD organization. Altogether, our study reveals pervasive regulatory function of RBPs in 3D genome organization.

## Results

### TAD boundaries are hotspots for RBP binding

To explore the potential regulatory roles of RBPs in 3D genome organization, we harnessed the ChIP-seq data performed on a large number of RBPs (44 in K562 and 28 in HepG2 cells) [20], together with Hi-C, RNA-seq, GRO-seq, and CLIP-seq (Suppl. Table S1) for integrated analyses as outlined in Fig. 1A. By processing the Hi-C data [29, 30], we identified a total of 10803 and 10958 TAD boundaries in K562 and HepG2, respectively (Suppl. Table S2). Overlaying with each of the 44 RBP ChIP-seq dataset in K562, we found a portion (ranging from 7.7 to 24.7%) of each RBP binding peaks overlapped with the above defined TAD boundaries and the remaining with intra-TAD regions (Fig. 1B), and TAF15 exhibited the highest (24.6%) enrichment on the TAD boundaries, which was close to the levels of cohesin subunits (RAD21, 29.3% andSMC3, 25.9%). In HepG2, 5.2% to 25.6 % of the RBP binding peaks were located on the boundaries with TARDBP ranking on the top (25.6%) (Fig. 1C). Furthermore, we found that a total of 48.1% (5198/10803) and 40% (4385/10958) of boundaries contained at least one RBP binding site in K562 and HepG2, respectively (Suppl. Fig. S1A).

**Figure 1.**
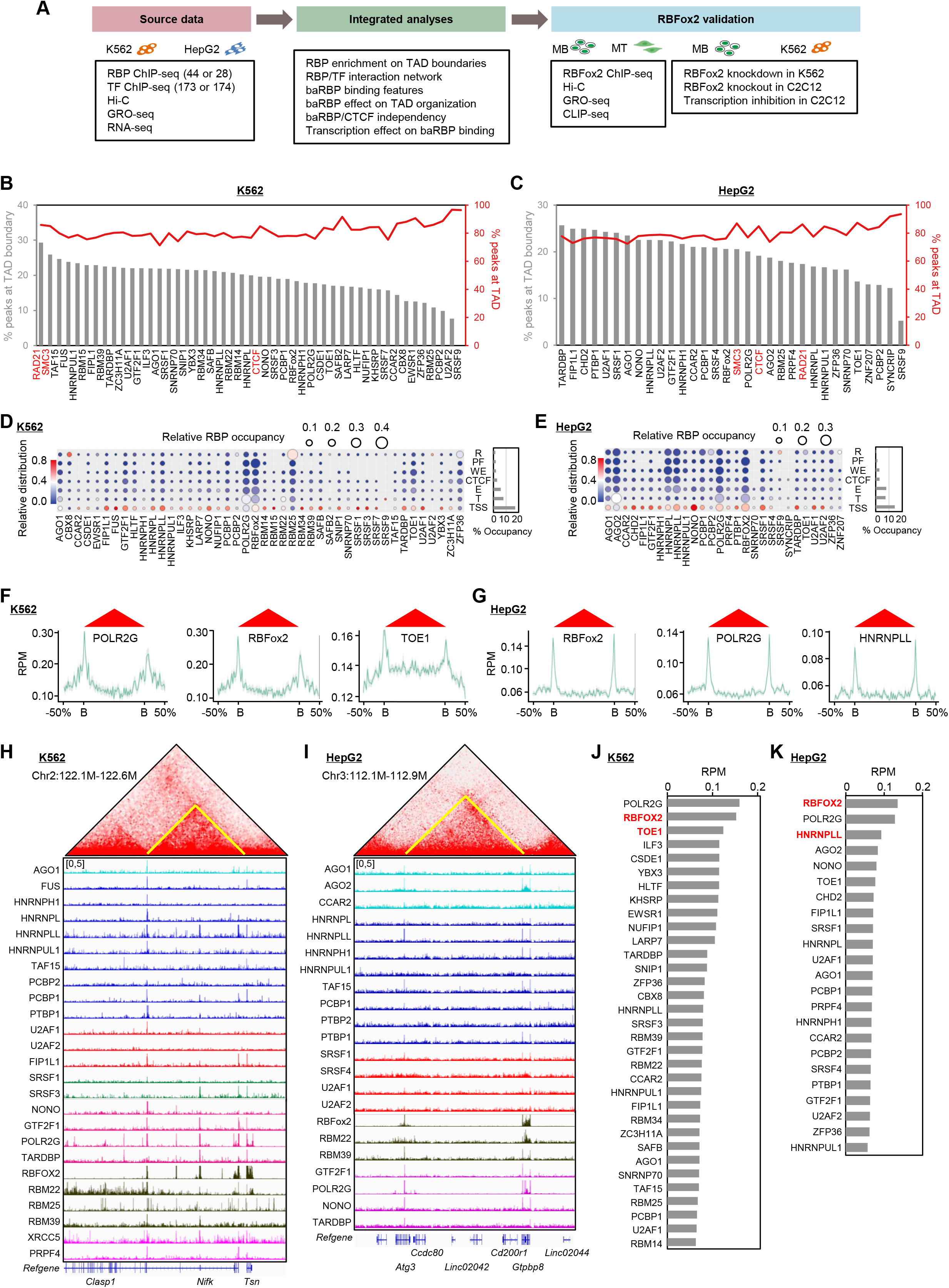
**TAD boundaries are hotspots for RBP binding. (A**) Schematic illustration of the analysis pipeline. Various datasets were collected from both K562 and HepG2 cells and integrated to elucidate the role of RBPs in 3D genome organization. **(B-C)** Bar plot showing the percentages of RBP ChIP-seq peaks overlapping with TAD boundaries (left y-axis) or TADs (right y-axis) in K562 and HepG2 cells. CTCF, cohesin subunit RAD21 and SMC3 ChIP-seq peaks are served as positive controls. **(D-E)** RBP occupancy on specific states of ENCODE-annotated genome segmentation of TAD boundaries in K562 and HepG2 cells. R, repressive regions; PF, promoter flanking regions; T, transcribed regions; CTCF, CTCF-binding sites; WE, weak enhancers; E, enhancers; TSSs, transcription start sites/promoters. For each RBP, the relative distribution of its occupied sites on individual segments (vertical comparison) is color coded (key on the left). For each class of segment annotation, the relative distribution of individual RBPs is represented by bubble size (horizontal comparison), as indicated by relative RBP occupancy. Right:summed percentage of individual segment annotations covered by the surveyed RBPs. **(F-G)** Meta-gene plots of representative RBPs (POLR2G, RBFox2, TOE1 in K562 and RBFox2, POLR2G and HNRNPLL in HepG2 cells) ChIP-seq signals around TAD boundaries. B:boundary. **(H-I)** Representative heatmaps showing the enrichment of RBP ChIP-seq signals on TAD boundaries in K562 and HepG2 cells. **(J-K)** Ranking of RBP ChIP-seq signals at TAD boundaries in K562 and HepG2 cells.

To further characterize the binding preference of these RBPs on TAD boundaries, we assigned ChIP-seq peaks of each RBP to seven ENCODE annotated genome segmentations, TSS:transcriptional start site; T:transcribed; R:repressed; PF:promoter flanking region; E:enhancer; WE:weak enhancer; CTCF binding region, and assessed the relative distribution of peaks among these segmentations (Fig. 1D-E). We found that TSSs were hotspots for RBP binding (Fig. 1D-E, vertical comparison). Collectively, boundary locating RBPs occupied 17% of promoters in K562 and 14% in HepG2 cells. In addition, transcribed regions (8% in K562, 7.3% in HepG2), enhancer regions (5.8% in K562, 7.9% in HepG2) and CTCF (2.8% in K562, 2.8% in HepG2) were also actively occupied by RBPs, while only a small part of repressed regions (1.5% in K562, 0.5% in HepG2) and promoter flanking regions (1.9% in K562, 1.3% in HepG2) were occupied by RBPs. Further calculating the relative contribution of individual RBPs to each genomic segmentation type, we found that RBM25 and AGO2 were the dominant RBPs on repressive regions in K562 and HepG2, respectively. POLR2G and RBFox2 were enriched at transcribed regions in both cells and also displayed a preference for enhancer and promoter flanking regions. Furthermore, metagene analyses revealed that ChIP-seq signals of 75% RBPs (33 out of 44) in K562 (for example, POLR2G, RBFox2, and TOE1, shown in Fig. 1F) or 82% (23 out of 28) in HepG2 (for example, RBFox2, POLR2G and HNRNPLL, shown in Fig. 1G) were highly enriched on TAD boundaries when compared with intra-TAD regions (Fig. 1H-I and Suppl. Fig. S1B-C) and were henceforth defined as boundary associated RBPs (baRBPs). Upon ranking, POLR2G, RBFox2, and TOE1 were the top three in K562 (Fig. 1J) whilst RBFox2, POLR2G, and HNRNPLL ranked top in HepG2 (Fig. 1K). The subsequent analyses were thus exemplified on RBFox2 and TOE1 in K562 or RBFox2 and HNRNPLL in HepG2 considering POLR2G is not a classical RBP. Notably, 12 of the baRBPs were shared in both K562 and HepG2 (Suppl. Fig. S1D) and 7 out of the 33 baRBPs were experimentally validated chromatin-associated RBPs in K562 [31] (highlighted in Suppl. Fig. S1D).

To take a close look at baRBP binding features on TAD boundaries, we conducted motif enrichment analysis and found that many baRBP peaks contained GC-rich sequences (15 out of 33 in K562 and 17 out of 23 HepG2 cells) (Suppl. Fig. S1E-F). Of note, 2.5∼94% in K562 and 13.4∼80% in HepG2 of these baRBP peaks overlapped with CpG islands (Suppl. Fig. S1G-H). More interestingly, by leveraging the available DNA G-quadruplex (dG4) ChIP-seq data [32], we found that all baRBPs except for CBX8 in K562 overlapped with the dG4 peaks to 8∼51% in K562 and 11-48% in HepG2 cells (Suppl. Fig. S1I-J).

### Network interactions of baRBPs on TAD boundaries

Next, to examine possible synergistic action of baRBPs on TAD boundaries, we performed the co-association event analysis by non-negative matrix factorization (NMF) method (Fig. 2A). A unifying set of cis-regulatory elements (CREs) associated with individual RBPs were defined and an occupancy profile was constructed based on whether individual CREs were associated by each baRBP. In this analysis, it is necessary to pre-determine a reasonable factor number for groups. In our case, we found 5 and 4 as an optimal group number in K562 and HepG2, respectively (Fig. 2B-C), with which the clustering became stabilized as revealed by cophenetic and dispersion correlation coefficients (Suppl. Fig. S2A-B) and the consensus matrix (Suppl. Fig. S2C-D). As a result, we found that POLR2G, RBFox2, AGO1, and SRSF9 were tightly clustered together as group 3 in K562 (Fig. 2B, D, Suppl. Table S3), while PCBP2, ZNF207, SYNCRIP, SRSF9, ZFP36, HNRNPLL, TOE1, and HNRNPUL1 were co-localized as group 3 in HepG2 (Fig. 2C, E, Suppl. Table S3). Notably, multiple factors in the identified groups were previously known to have physical interactions at protein level [33] (blue edges in Fig. 2D, E).

**Figure 2.**
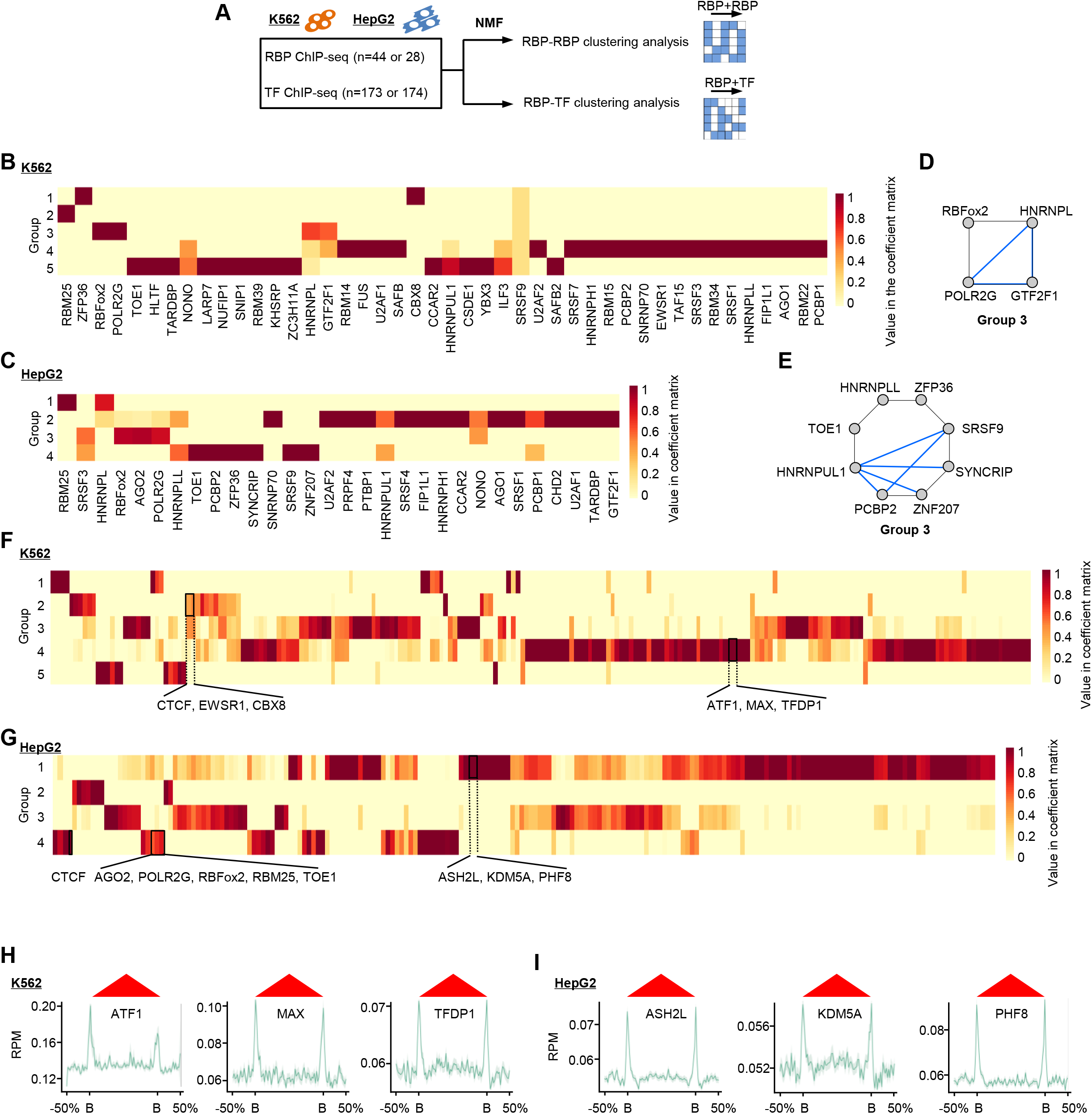
Network interaction of baRBPs and TFs at TAD boundaries. (A) Schematic illustration of the analysis pipeline. baRBP and TF ChIP-seq dataset were collected from both K562 and HepG2 cells and integrated to perform clustering analysis by using NMF method. **(B-C)** Segregation of baRBP peaks residing at TAD boundaries into 5 and 4 groups by NMF-inferred coefficient matrixes in K562 and HepG2 cells. **(D-E)** Representative NMF-segregated groups in K562 and HepG2 cells. Blue line:known physical interactions between members in each group annotated by GeneMANIA. **(F-G)** Segregation of baRBPs and TF peaks residing at boundaries into 5 and 4 groups by NMF-inferred coefficient matrixes in K562 and HepG2 cells. **(H-I)** Meta-gene plot of the ChIP-seq signals around TAD boundaries of three representative TFs in K562 (ATF1, MAX, TFDP1) and HepG2 cells (ASH2L, KDM5A, PHF8). B:boundary.

Furthermore, to test if components of active transcription complexes such as transcription pre-initiation complex (PIC) or TFs may partner with RBPs on TAD boundaries, we took advantage of the rich resources of available TF ChIP-seq datasets (173 TFs in K562 and 174 TFs in HepG2) (Fig. 2A); By performing NMF analysis (Suppl. Fig. S2E-H), four groups of baRBPs/TFs were identified in K562 and five in HepG2. Of note, structure protein such as CTCF co-localized with baRBPs such as CBX8 and EWSR1 in K562 (Fig. 2F), and CTCF and cohesin co-localized with AGO2, POLR2G, RBFox2, RBM25, and TOE1 in HepG2 (Fig. 2G). Interestingly, meta-gene analyses showed that some TFs such as ATF1, MAX, and TFDP1 in K562 (Fig. 2H) and ASH2L, KDM5A, and PHF8 in HepG2, were also enriched on TAD boundaries (Fig. 2I). Altogether, these results revealed potential synergistic functions of baRBPs and also with TFs in regulating TAD organization.

### baRBP enrichment correlates with increased insulation strength of TAD boundaries

To further investigate the possible effects of each baRBP binding on TAD organization, we performed Aggregate Domain Analysis (ADA) and found that the intra-TAD interactions of the TADs with baRBP bound (+baRBP) boundaries were much higher compared with those without baRBP (-baRBP) boundaries in K562 (Fig. 3A, Suppl. Table S4). Consistently, we found that boundaries with each of the 44 baRBPs (+baRBP) displayed significantly lower insulation score thus higher insulation strength compared to those without baRBPs (-baRBP) (Suppl. Fig. S3A and Suppl. Table S4). For example, RBFox2 and TOE binding correlates with 1.35 and 1.43-fold increase of insulation score (IS) (Fig. 3B); domain score, which quantifies a TAD’s tendency to self-interact, was also higher on the TADs with +baRBP boundaries (Fig. 3C, Suppl. Table S4). These findings were further substantiated by the results from 2D pile-up contact maps around boundaries showing that +baRBP boundaries exhibited higher intra-and inter-TAD interaction frequency (Fig. 3D-E) compared with those-baRBP boundaries. Similar findings were found in HepG2 cells except for the inter-TAD interaction frequency was lower on +baRBP boundaries (Fig. 3F-J, Suppl. Fig. S3B and Suppl. Table S4); binding of each of the 23 baRBPs was correlated with a much higher IS (Suppl. Fig. S3B and Suppl. Table S4), which was exemplified by RBFox2 and HNRNPLL (1.32 and 1.37-fold increase of IS, respectively) (Fig. 3G). Altogether, the above findings suggest that baRBPs may have a positive effect on TAD insulation. Nevertheless, we found no significant association between the number of baRBP binding sites with insulation score; for example, the number of RBFox2 binding peaks at boundaries was not associated with insulation score (Supp. Fig. S4A, left and middle panels), and the RBFox2 ChIP-seq signals were not associated with insulation score (Suppl. Fig. S4A, right panel). Further analyses demonstrated that the effects of baRBP binding on TAD boundaries (Suppl. Fig. S4B-G) could be observed in both A and B compartments. Nevertheless, the impact of the binding on intra-TAD interactions was more pronounced in B vs A compartment (Suppl. Fig. S4B-E). Altogether, the above results demonstrate that baRBP enrichment on TAD boundaries is associated with enhanced insulation ability thus may facilitate TAD organization. The above findings are illustrated on several loci on chromosome 1 (Fig. 3K-L). For example, in K562, a TAD encompassing *Spocd1* and*Ptp4a2* genes with-RBFox2 boundaries (Fig. 3K, left panel) displayed a much lower domain score than a TAD with +RBFox2 boundaries (Fig. 3K, right panel). Similarly, in HepG2, a TAD encompassing *Camta1* gene with-HNRNPLL boundaries (Fig. 3L, left panel) displayed a lower domain score than a TAD with +HNRNPLL boundaries (Fig. 3L, right panel).

**Figure 3.**
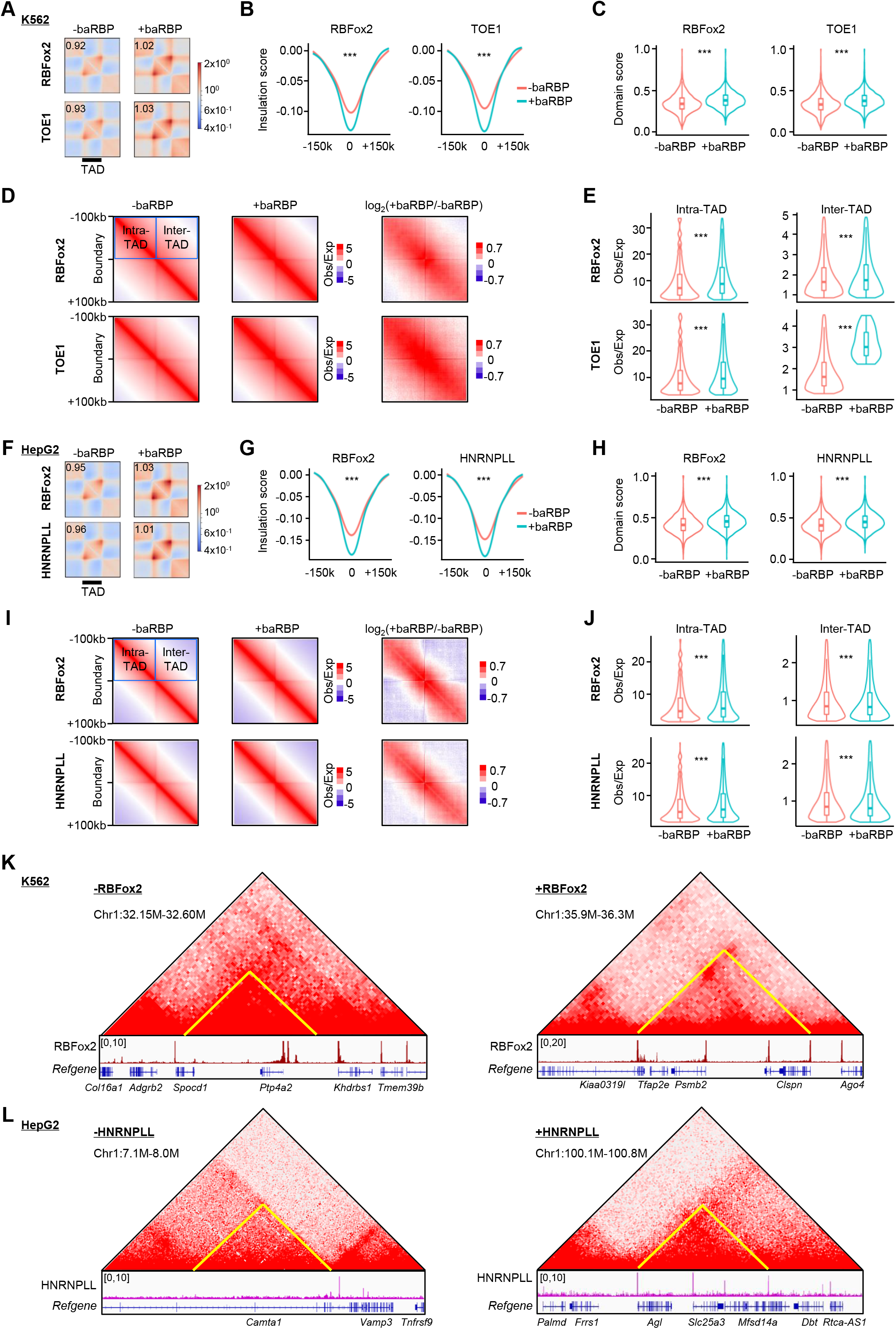
baRBP enrichment correlates with increased insulation strength of TAD boundaries (A) Aggregate domain analysis (ADA) of TADs with (+) or without (-) representative baRBPs (RBFox2 and TOE1) boundaries in K562 cells. **(B)** Comparison of insulation scores of +baRBP and-baRBP boundaries. **(C)** Comparison of domain scores of TADs with +baRBP or-baRBP boundaries in K562 cells. **(D)** Aggregate analysis of interaction frequency around +baRBP and-baRBP boundaries in K562 cells. **(E)** Quantification of interaction frequency around +baRBP and-baRBP boundaries in **(D)**. **(F-J)** The above analyses were performed in HepG2 cells for two representative baRBPs (RBFox2 and HNRNPLL and RBFox2). **(K-L)** Representative heatmaps and UCSC tracks showing the baRBP (RBFox2 in K562 and HNRNPLL in HepG2) ChIP-seq signals around TADs with +baRBP boundaries (left) or-baRBP boundaries (right).

It is also interesting to find out that TADs with +baRBP boundaries were significantly smaller in size compared to those without baRBPs (Suppl. Fig. S4H-I), and these TADs also encapsulated a significantly higher number of protein-coding genes (PCGs) per domain (Suppl. Fig. S4J-K).

### baRBPs may facilitate TAD organization independent of CTCF and cohesin binding

Next to examine the possible synergism between baRBPs and CTCF on TAD boundaries, we classified the boundaries into four groups based on the binding of CTCF and individual baRBP, and found that, for all baRBPs in K562, a large proportion of boundaries were bound by CTCF alone (+CTCF-baRBP) (ranging from 31% for RBFox2 to 65% for SAFB) while less than 10% of boundaries were bound by baRBP alone (-CTCF+baRBP) (Fig. 4A). Moreover, some baRBPs displayed a high co-occupancy with CTCF (+CTCF+baRBP) (for example, 34.4% of RBFox2) but others did not (0% for SAFB, SRSF9, and U2AF2) (Fig. 4A). Interestingly, a rather high portion of boundaries (ranging from 25-35%) were negative for CTCF and baRBP binding (-CTCF-baRBP). Similar observations were made in HepG2 cells:the proportion of +CTCF-baRBP boundaries ranged from 30% (AGO2) to 65% (SRSF9)), while less than 17% of boundaries were-CTCF+baRBP (Fig. 4B). The percentage of +CTCF+baRBP boundaries ranged from 0% (SRSF9) to 35.4% (AGO2) (Fig. 4B). Moreover, when we took a closer examination of the CTCF and baRBP binding peaks at the +CTCF+baRBP boundaries, the overlapping ranged from 20% to 30% (Fig. 4C-D, Suppl. Table S5), indicating a potential synergism of baRBPs and CTCF on TAD organization. Nevertheless, on CTCF-present boundaries, further analyses uncovered that baRBP binding was not associated with increased insulation strength (Fig. 4E, Suppl. Fig. S5A-B); and increased intra-TAD interaction frequency was observed in HepG2 but not in K562 (Fig. 4F, Suppl. Table S4). The baRBP binding also showed an association with higher domain score in both cells (Fig. 4G). However, on CTCF-absent boundaries, baRBP binding was associated with significantly higher insulation strength (Fig. 4H, Suppl. Fig. S5C-D), intra-TAD interaction frequency (Fig. 4I) and domain score (Fig. 4J) in both K562 and HepG2 cells. As illustrated in Fig. 4K-L, a TAD boundary encompassing *Pex14* gene, was co-bound by RBFox2 and CTCF and displayed higher insulation strength than a boundary with RBFox2 binding alone in K562 (Fig. 4K); similar phenomenon was illustrated in HepG2, a TAD boundary encompassing *Nek2* gene was co-bound by RBFox2 and CTCF and showed higher insulation strength than a boundary with RBFox2 binding alone (Fig. 4L). These results suggest that baRBP binding may exert the TAD regulatory function in a CTCF-independent manner.

**Figure 4.**
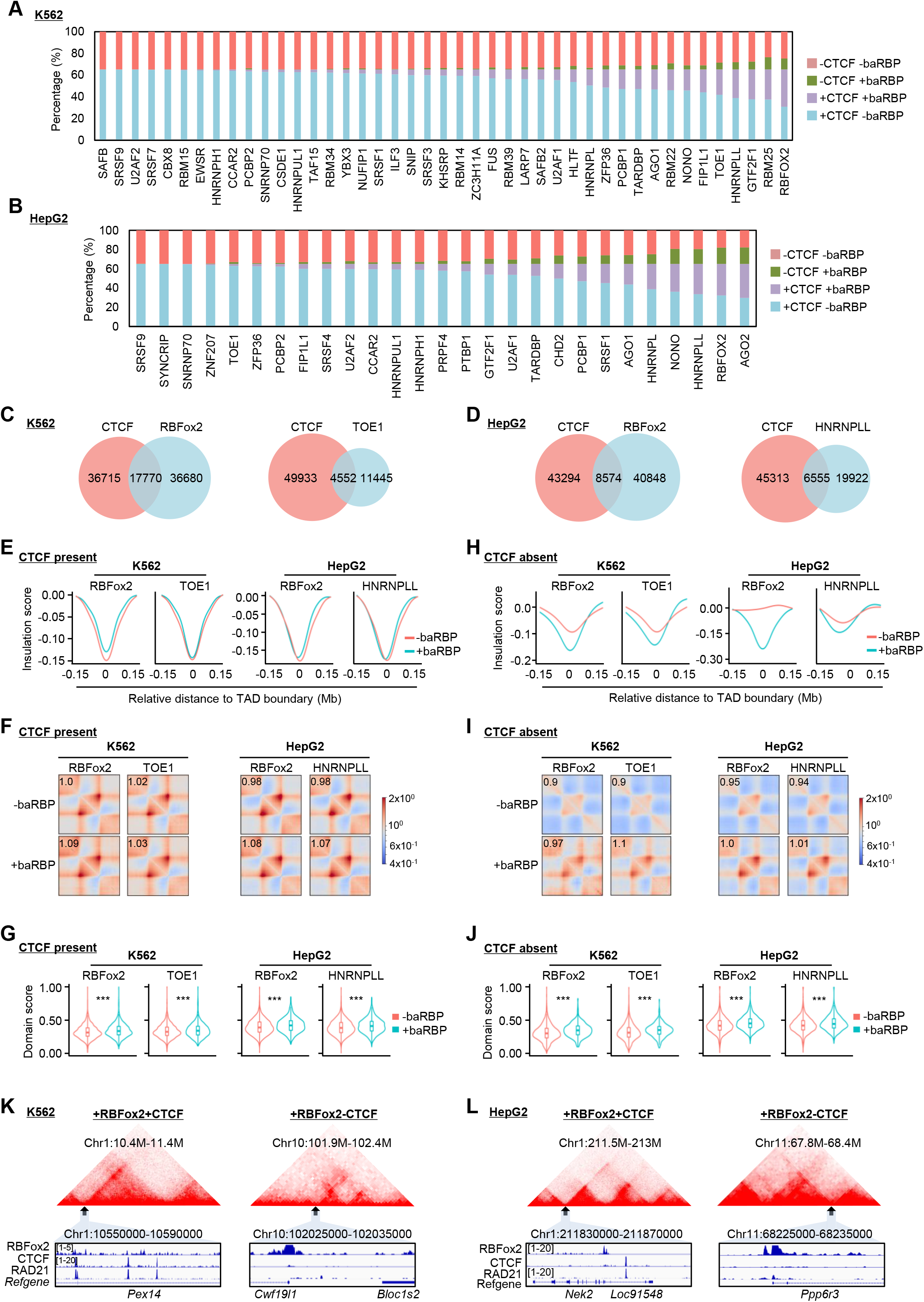
baRBPs may facilitate TAD organization independent of CTCF and cohesin binding. (A-B) Proportions of different types of boundaries in K562 and HepG2 cells. **(C-D)** Overlapping of CTCF and representative baRBPs (RBFox2 and TOE1 in K562; RBFox2 and HNRNPLL in HepG2) ChIP-seq peaks at boundary regions. **(E)** Comparison of insulation scores of +baRBP and-baRBP boundaries at CTCF present boundaries. **(F)** Aggregate domain analysis of TADs with or without baRBP at CTCF present boundaries. **(G)** Comparison of domain scores between TADs with or without baRBP at CTCF present boundaries. **(H-J)** The above analyses were performed at CTCF absent boundaries. **(K-L)** Representative heatmaps and UCSC tracks showing baRBP (RBFox2), CTCF, and cohesin ChIP-seq signals around CTCF present or CTCF absent boundaries in K562 and HepG2 cells.

To further examine possible synergism between baRBPs and cohesin on TAD organization considering physical interaction has been demonstrated between cohesin and several RBPs [34], we analyzed the binding of RAD21 and SMC3 and found both factors were highly enriched on +baRBP boundaries (Suppl. Fig. S6A-B). Moreover, 10%-20% of the baRBP peaks overlapped with RAD21 peaks on the boundaries (Suppl. Fig. S6C-D, Suppl. Table S5). Similar to CTCF, we found that on RAD21 present boundaries, baRBP binding was not associated with insulation strength (Suppl. Fig. S6E-F), but significantly associated with increased intra-TAD interactions (Suppl. Fig. S6G) and domain scores (Suppl. Fig. S6H). On RAD21 absent boundaries, on the other hand, baRBP binding was associated with increased insulation strength (Suppl. Fig. S6I-J), intra-TAD interaction frequency (Suppl. Fig. S6K) and domain score (Suppl. Fig. S6L). For example, TAD boundaries that were co-bound by RBFox2 and RAD21 displayed higher insulation strength than those bound by RBFox2 alone in K562 (Fig. 4K-L). Altogether, the above findings suggest that baRBP binding may facilitate TAD organization independent of CTCF or cohesin.

### baRBP enrichment on TAD boundaries is correlated with active transcription

Increasing evidence shows that RBPs can be recruited to chromatin by nascent RNAs or eRNAs [23, 27, 35, 36]. To reveal the possible connection between active transcription and baRBP binding, we analyzed the available GRO-seq datasets from K562 and HepG2 and found that in both cells, +baRBP boundaries were indeed associated with much higher nascent transcription levels compared to-baRBP boundaries (Fig. 5A, Suppl. Fig. S7A-B, Suppl. Table S6). When we further examined the transcriptional levels of random regions (shuffled) and baRBP binding regions within the boundaries, we also observed significantly higher transcriptional levels at the +baRBP regions (Fig. 5B). Moreover, the level (low, medium or high) of baRBP ChIP-seq signal was positively correlated with the GRO-seq signals (Fig. 5C). It has been reported that active transcription facilitates the formation of TAD or the establishment of insulation ability of boundaries[16, 37, 38]; inhibiting transcription initiation or elongation decreases TAD boundary strength and loop interaction[38, 39]. Indeed, we found that the majority of +baRBPs boundaries (29 out of 33 in K562, 12 out of 23 in HepG2) with nascent transcription displayed significantly higher insulation strengh compared to those without transcription (Fig. 5D and Suppl. Fig. S7C-F), and the level of GRO-seq signals (low, medium, or high) was negatively correlated with the insulation score (Fig. 5E and Suppl. Fig. S7G-H). Moreover, the transcriptional level was also positively correlated with the intra-TAD interaction frequency (Suppl. Fig. S7I-L), which was supported by representative Hi-C contact heatmaps (Fig. 5F). Altogether, the above findings demonstrate that baRBP enrichment on TAD boundaries may be facilitated by active transcription and baRBP/transcription may synergistically promote TAD boundary insulation.

**Figure 5.**
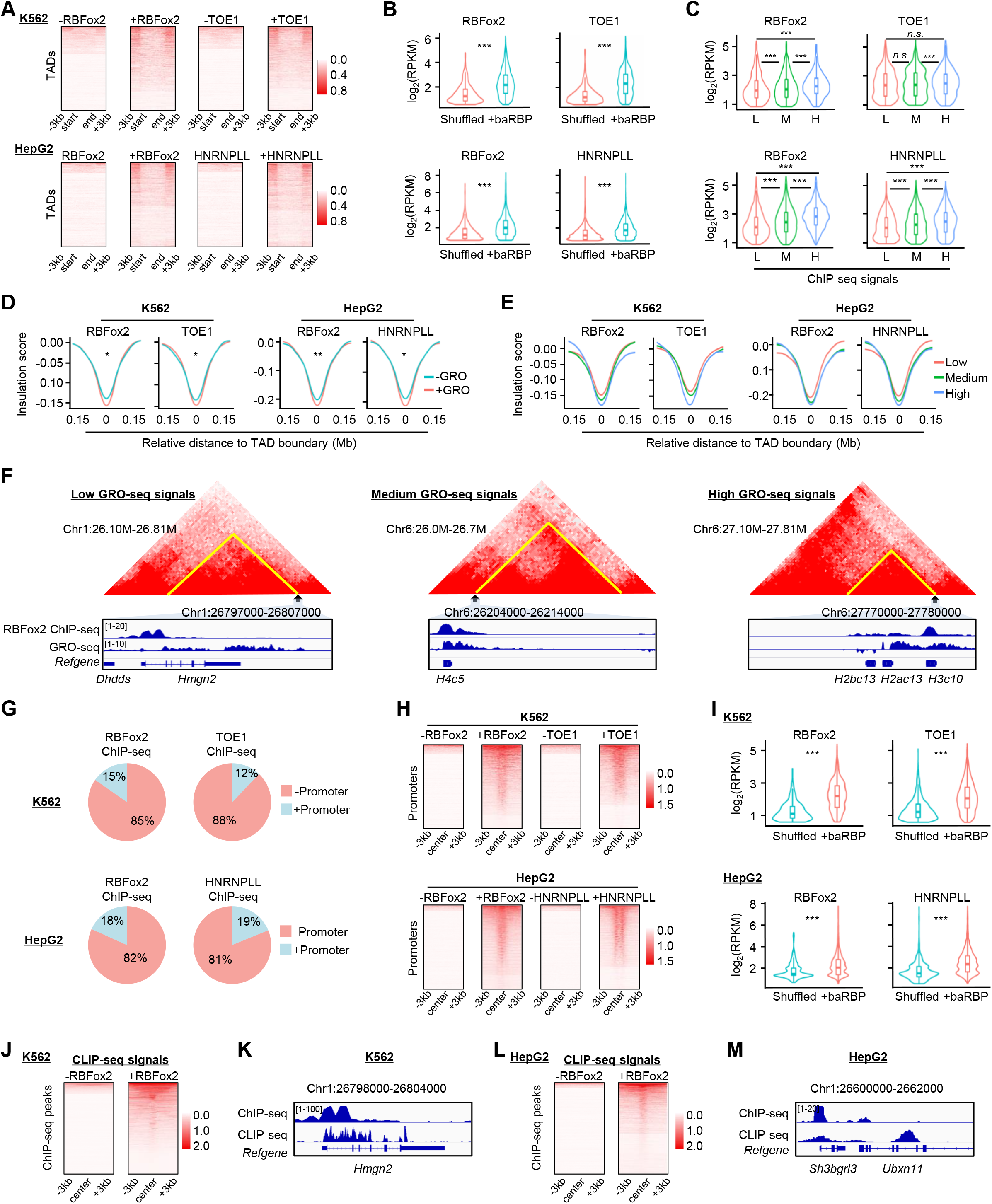
baRBP enrichment at TAD boundaries is correlated with active transcription. (A) Heatmaps showing the GRO-seq signals around TADs with (+) or without (-) baRBP (RBFox2 and TOE1 in K562, RBFox2 and HNRNPLL in HepG2) boundaries. **(B)** Comparison of GRO-seq signals at +baRBP and random regions (shuffled) in K562 and HepG2 cells. **(C)** Comparison of GRO-seq signals at TAD boundaries with low (L), medium (M) or high (H) levels of +baRBP signals in K562 and HepG2 cells. **(D)** Comparison of insulation scores of +baRBP boundaries with (+) or without (-) GRO-seq signals in K562 and HepG2 cells. **(E)** Comparison of insulation scores of +baRBP boundaries with different levels of GRO-seq signals in K562 and HepG2 cells. **(F)** Representative heatmaps and tracks showing TADs with +baRBP (RBFox2) boundaries with different levels of GRO-seq signals in K562 cells. **(G)** Pie charts showing the proportions of baRBP (RBFox2 and TOE1 in K562, RBFox2 and HNRNPLL in HepG2) ChIP-seq peaks residing at promoter containing boundaries (+promoter boundaries). **(H)** Heatmap showing the distribution of GRO-seq signals around +baRBP (RBFox2 and TOE1 in K562, RBFox2 and HNRNPLL in HepG2) or randomly shuffled regions at +promoter boundaries. **(I)** Comparison of GRO-seq signals of baRBP binding sites and random region at +promoter boundaries in K562 and HepG2 cells. **(J)** Heatmaps showing the distribution of RBFox2 CLIP-seq signals around baRBP binding sites (RBFox2 in K562) or randomly shuffled region. **(K)** Representative tracks showing the overlapping of RBFox2 ChIP-seq and CLIP-seq signals in K562 cells. **(L-M)** The above analyses were performed on RBFox2 in HepG2 cells.

Since promoter transcription is known to facilitate TAD boundary formation [7], we next investigated if baRBP binding on TAD boundaries is mediated through active promoter transcription. Indeed, a portion (5.4% to 20.2% in K562 and 7.6% to 22.5% in HepG2, Suppl. Table S7) of baRBP ChIP-seq peaks were located at promoter-containing boundaries (+promoter boundaries) (Fig. 5G) despite only a small portion (∼30%) of TAD boundaries contained gene promoters (data not shown). As expected, on +promoter boundaries, GRO-seq signals at the +baRBP binding sites were much higher than those random regions (shuffled) (Fig. 5H-I). Moreover, by incorporating the available CLIP-seq data for RBFox2 in both K562 and HepG2 cells, we found that RBFox2-bound RNA signals were enriched around its DNA binding regions (Fig. 5J-M), suggesting that RBFox2 possibly binds with promoter transcripts co-transcriptionally. Additionally, since it is reported that eRNAs can mediate AGO1 regulation of 3D genome organization via interacting with AGO1 and chromatin [27], we also investigated the possibility of baRBP binding with eRNAs and found no such evidence (data not shown). Altogether, the above findings demonstrate that co-transcriptional binding of baRBPs with promoter transcripts may play an active role in TAD organization.

### RBFox2 binding promotes TAD organization in K562

To further illuminate the causative role of baRBPs in TAD organization, we decided to select RBFox2 as a paradigm for in-depth experimental dissection considering its top ranking in all the above analyses in both K562 and HepG2. RBFox2 is known as a key regulator of alternative exon splicing in cells[35, 40, 41] but recent evidence unveiled that RBFox2 interacts with chromatin via nascent RNAs to mediate transcriptional repression [35]. To this end, we knocked down *RBFox2* in K562 using siRNA oligos and examined its effect on 3D organization (Fig. 6A-B). We found that *RBFox2* knock-down had limited effect on global 3D genome organization as no difference in the contact frequency was detected between *siNC* and *siRBFox2* at both short-and long-distance levels (Suppl. Fig. S8A). The knock-down did not alter the overall genome organization at compartment level either (Suppl. Fig. S8B, Suppl. Table S8), with only 2% of compartments switched either from A to B or B to A, and the percentage of RBFox2 binding had no significant difference at the switched compartments (Suppl. Fig. S8C-D). Nevertheless, to quantify the compartmental interactions, we generated saddle plots and found a significant increase in top 20% B-B but not top 20% A-A compartmental interactions upon *RBFox2* deletion (Suppl. Fig. S8E-F). At the TAD level, we did not observe significant changes on the number (Suppl. Fig. S8G, Suppl. Table S8) and size (Suppl. Fig. S8H) upon *RBFox2* deletion. TAD score remained unaltered (Suppl. Fig. S8I) and the intra-TAD interactions as measured by ADA was also not affected (Suppl. Fig. S8J). By performing APA, we also did not detect significant change of loop interactions (Suppl. Fig. S8K). Nevertheless, a global re-organization of TAD boundaries was observed upon *RBFox2* loss; A total of 4718 boundaries were identified in *siNC* cells and a large portion (57%) were lost upon *RBFox2* depletion (Fig. 6C). Still, the overall insulation score remained unaltered (Fig. 6D, Suppl. Table S8).

**Figure 6.**
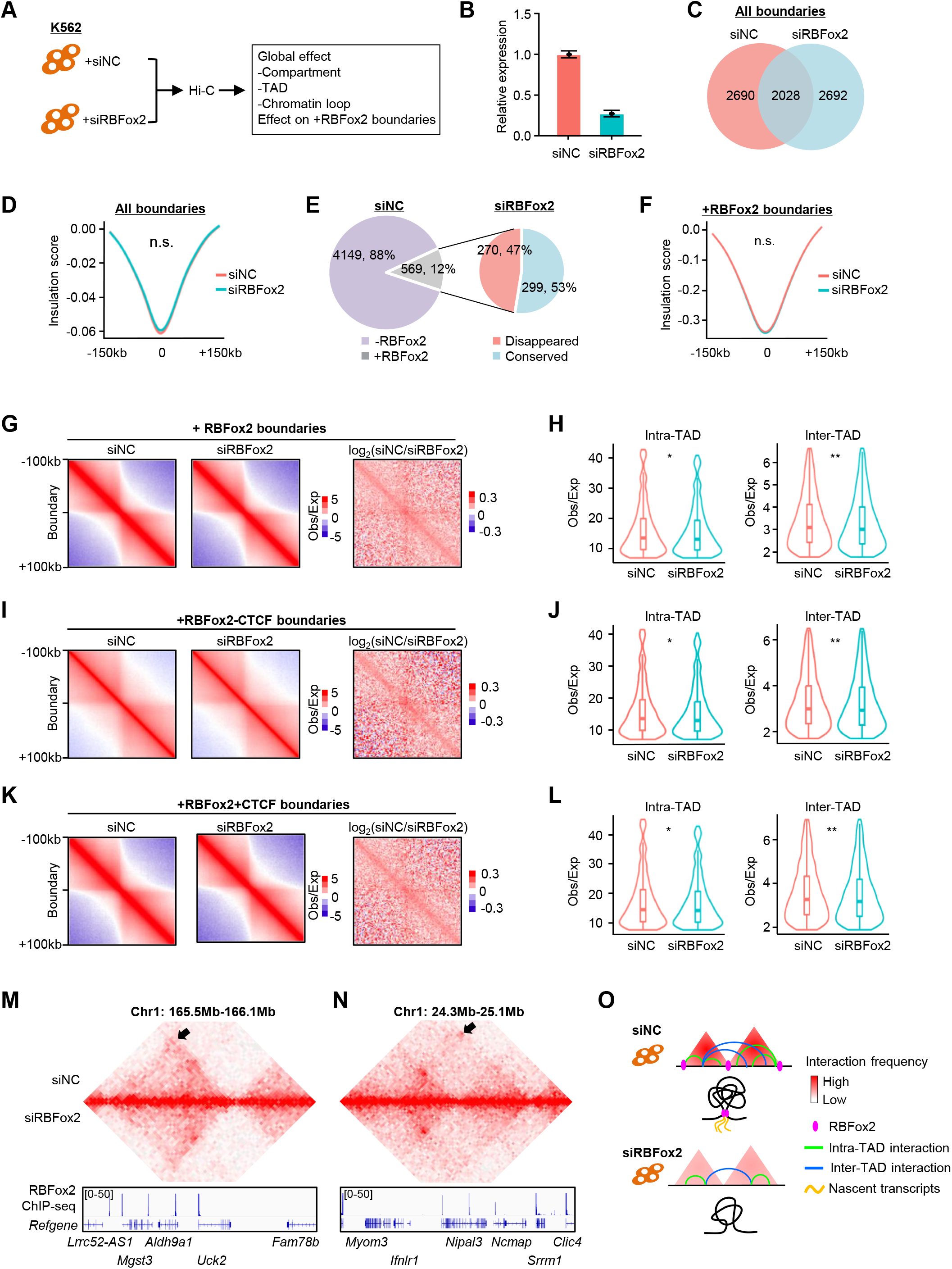
RBFox2 binding promotes TAD organization in K562. (A) Schematic illustration of the experimental testing of *RBFox2* knockdown effect on 3D genome in K562 cells. **(B)** *RBFox2* knockdown efficiency was detected by RT-qPCR. **(C)** Dynamic changes of all boundaries upon *RBFox2* depletion. **(D)** Comparison of insulation scores of all boundaries in *siNC* and *siRBFox2* cells. **(E)** Percentage of +RBFox2 boundaries in *siNC* sample (left) and the percentage of disappeared or conserved boundaries in *siRBFox2* cells (right). **(F)** Comparison of insulation scores of +RBFox2 boundaries in *siNC* and *siRBFox2* cells. **(G)** Aggregate analysis of interactions around +RBFox2 boundaries. **(H)** Comparison of inter-and intra-TAD interactions around +RBFox2 boundaries in *siNC* and *siRBFox2* cells. **(I)** Aggregate analysis of interactions around +RBFox2-CTCF boundaries in *siNC* and *siRBFox2* cells. **(J)** Comparison of inter-and intra-TAD interactions around +RBFox2-CTCF boundaries in *siNC* and *siRBFox2* cells. **(K-L)** The above analyses were performed on +RBFox2+CTCF boundaries. **(M-N)** Two representative heatmaps displaying the decreased intra-TAD interactions upon *RBFox2* depletion. **(O)** Schematic illustration of the role of RBFox2 on TAD boundary organization in K562 cells.

We then took a close examination at the TADs with +RBFox2 boundaries in *siNC* and observed notable reorganization with 294, 309, 3, and 108 TADs merged, split, disappeared, or rearranged (Suppl. Fig. S8L). Moreover, 47% of the 569 +RBFox2 boundaries disappeared upon *RBFox2* deletion (Fig. 6E), indicating the importance of RBFox2 binding in maintaining these boundaries. Nevertheless, *RBFox2* loss did not have significant impact on insulation scores of these boundaries (Fig. 6F) as both intra-and inter-TAD interactions were decreased simultaneously (Fig. 6G-H). When further examining the +RBFox2-CTCF boundaries, similar phenomenon was observed with no significant impact on insulation score due to simultaneous attenuation of both intra-and inter-TAD interactions (Fig. 6I-J). And this was also observed on +RBFox2+CTCF boundaries (Fig. 6K-L). Two representative interaction heatmaps on chr1 shown in Fig. 6M-N further illustrate the decreased intra-TAD interactions upon *RBFox2* depletion (Fig. 6M-N). Altogether, the above findings demonstrate a potential causative effect of RBFox2 binding on TAD organization (Fig. 6O).

### RBFox2 regulation of 3D genome is relevant in mouse myoblast differentiation

To expand our findings beyond K562 and HepG2 cells, we next examined the potential TAD regulatory function of RBFox2 in C2C12 myoblast (MB) cells by leveraging various datasets generated from our group and others (Fig. 7A, Suppl. Table S1) [24, 42, 43]. MBs can undergo differentiation into myotubes (MTs) which is an important process for skeletal muscle development and regeneration [44] to repair damaged muscles and RBFox2 expression was increased during the differentiation process (Fig. 7B) as previously reported [40]. First, by interrogating the Hi-C and RBFox2 ChIP-seq data from MBs (Suppl. Table S9), we found that RBFox2 binding signals were highly enriched on TAD boundaries (Fig. 7C), which is consistent with findings from K562 and HepG2 cells (Fig. 1). Similarly, +RBFox2 boundaries (11182) showed a much higher insulation strength compared with those without its binding (Fig. 7D). Consequently, genes within these TADs also displayed higher expression levels (Fig. 7E), and these genes were enriched for GO terms related with RNA processing (Suppl. Fig. S9A). Furthermore, we found that +RBFox2 boundaries underwent pronounced remodeling when MBs differentiated to MTs; 44.5% were lost (Fig. 7F) and increased insulation strength was observed (Fig. 7G). For example, the TAD encompassing *Au021092* gene showed increased intra-TAD interaction upon the differentiation from MB to MT (Fig. 7H). These observations suggest that RBFox2 may play an active role in boundary organization which is relevant to myoblast differentiation process.

**Figure 7.**
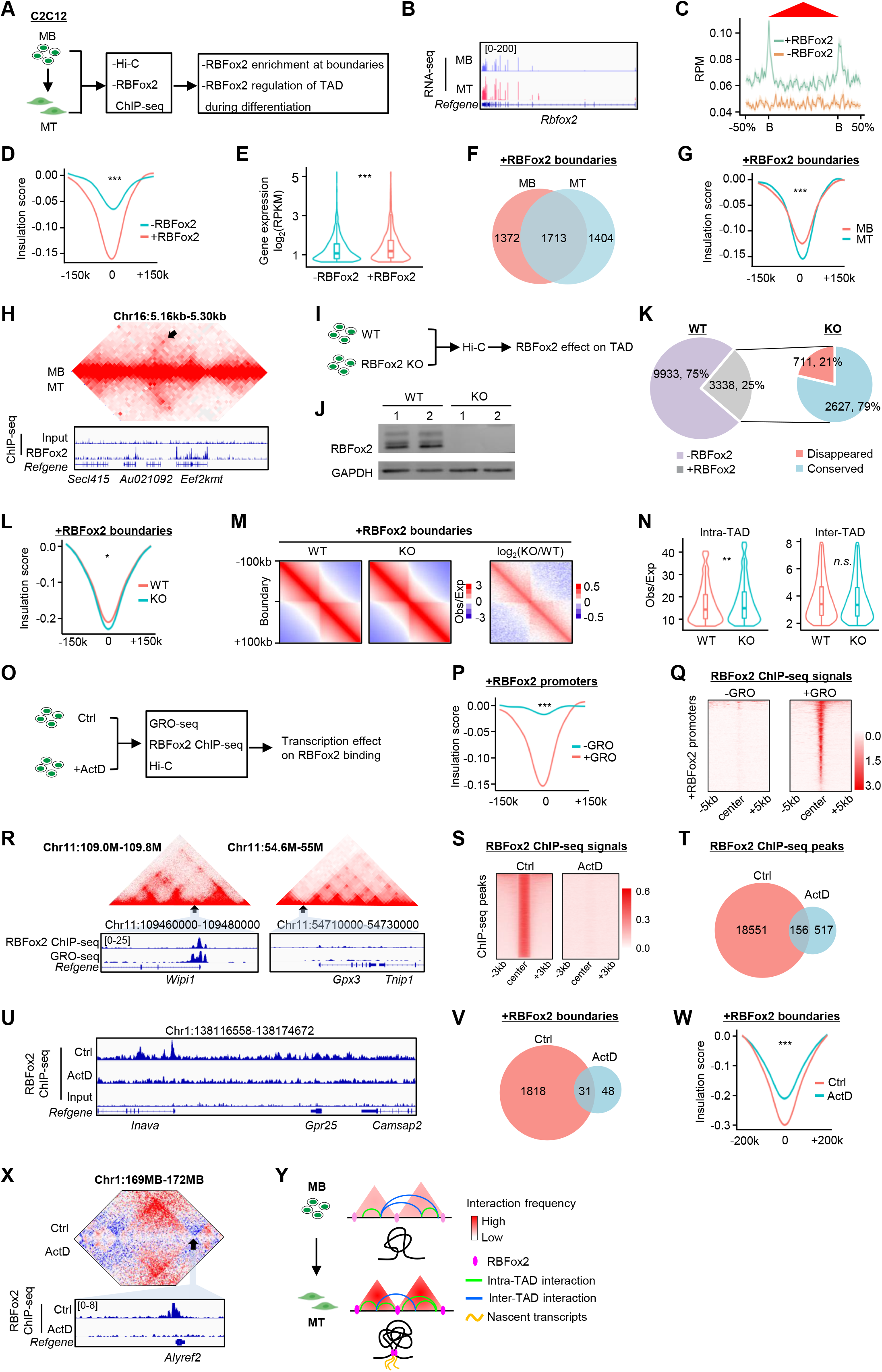
RBFox2 regulation of 3D genome is relevant in mouse myoblast differentiation. (A) Schematic illustration of experimental testing of RBFox2 regulation of 3D genome during C2C12 myoblasts (MBs) differentiation into myotubes (MTs). **(B)** UCSC track showing the expression of RBFox2 in MB and MT. **(C)** Meta-gene analysis showing the enrichment of RBFox2 ChIP-seq signals at TAD boundaries in MBs. **(D)** Comparison of insulation scores of +RBFox2 and-RBFox2 boundaries in MBs. **(E)** Comparison of expression of genes residing in TADs with +RBFox2 or-RBFox2 boundaries. (**F)** Dynamic changes of +RBFox2 boundaries during MB to MT differentiation. **(G)** Comparison of insulation scores of +RBFox2 boundaries in MB vs MT. **(H)** Representative heatmap displaying the increased interaction around +RBFox2 boundaries in MT vs MB. **(I)** Schematic illustration of the experimental testing of *Rbfox2* knockout (KO) effect on 3D genome in MBs. **(J)** Western blot detection of the knockout of RBFox2 protein in MBs by *sgRNAs* oligos. **(K)** Percentage of +RBFox2 boundaries in WT (left) and the percentage of disappeared or conserved boundaries in KO cells (right). **(L)** Comparison of insulation score of +RBFox2 boundaries upon *Rbfox2* KO. **(M)** Aggregate analysis of interactions around +RBFox2 boundaries. **(N)** Comparison of inter-and intra-TAD interactions around +RBFox2 boundaries in **(M)**. **(O)** Schematic illustration of the experimental testing of transcription effect on RBFox2 binding in MBs. **(P)** Comparison of insulation scores of +RBFox2 boundaries with or without GRO-seq signals in MB. **(Q)** Heatmaps showing RBFox2 ChIP-seq signals at +GRO-seq and-GRO-seq sites. **(R)** Two representative heatmaps displaying the interaction and RBFox2 ChIP-seq signals at +GRO-seq and-GRO-seq sites. **(S)** Heatmaps showing RBFox2 ChIP-seq signals on chromatin in Ctrl or ActD treated MB. **(T)** Pie chart showing the decreased number of RBFox2 ChIP-seq peaks in ActD vs Ctrl MB. **(U)** Representative tracks showing the significant decreased RBFox2 binding upon ActD treatment. **(V)** Pie chart showing the decreased number of +RBFox2 boundaries in ActD vs Ctrl MB. **(W)** Comparison of insulation scores of +RBFox2 boundaries in ActD vs Ctrl MB. **(X)** Heatmap showing the decreased RBFox2 binding and decreased interaction at RBFox2 binding site upon ActD treatment. **(Y)** Schematic illustrating the role of RBFox2 on TAD organization during C2C12 MB differentiation into MT.

To strengthen the above results, we knocked out *Rbfox2* (KO) in C2C12 MBs (Fig. 7I-J). Similar to finding from K562 cells, *Rbfox2* KO caused no detectable effect on global 3D genome and compartment organization (Suppl. Fig. S9B-G). TAD boundaries underwent dynamic changes (Suppl. Fig. S9H, Suppl. Table S10), but the insulation strength of all TAD boundaries and inter-TAD interaction frequency were unchanged (Suppl. Fig. S9I-K, Suppl. Table S10). When we further examined the 3338 +RBFox2 boundaries in WT, we found 21% of these boundaries disappeared upon *Rbfox2* KO (Fig. 7K). Nevertheless, RBFox2 bound boundaries showed a slightly increased insulation strength (Fig. 7L), which was opposite to the finding from K562 cells (Fig. 6), and the intra-TAD interaction frequency was significantly strengthened (Fig. 7M-N). For example, the boundary encompassing *Fam8a1* gene displayed an increased insulation strength upon *Rbfox2* KO, similar finding was also made on the boundary encompassing *Lipg* gene (Suppl. Fig. S9L-M)

Furthermore, to examine if RBFox2 association with TAD boundaries in C2C12 cells is mediated through nascent transcription (Fig. 7O), we found that 17% of RBFox2 binding peaks resided in active promoters (data not shown); and these +RBFox2 promoters with active GRO-seq signals showed a markedly higher insulation strength (Fig. 7P) and RBFox2 binding (Fig. 7Q) compared to those without GRO-seq signals. As illustrated in Fig. 7R, the promoter of *Wipi1* but not *Gpx3* was highly enriched for RBFox2 ChIP-seq and GRO-seq signals. To strengthen the findings, we treated C2C12 MBs with a transcriptional inhibitor Actinomycin D (ActD) and found the treatment attenuated the interaction frequency at short-range (1-40 Kb) but increased it at long-range (1-10 Mb) compared with non-treated control (Ctrl) cells (Suppl. Fig. S9N). At the TAD level, we observed the dynamic changes of TAD boundaries upon ActD treatment (Suppl. Fig. S9O, Suppl. Table S11). The intra-TAD interactions measured by ADA showed no alteration upon the treatment (Suppl. Fig. S9P) but the insulation strength and intra-TAD interactions of all boundaries were attenuated (Suppl. Fig. S9Q-S), supporting a role of transcription in boundary formation. Expectedly, by performing RBFox2 ChIP-seq in the above cells, we found that ActD treatment decreased the global RBFox2 binding signals (Fig. 7S-U) and reduced the number of +RBFox2 boundaries (Fig. 7V); insulation and intra-TAD interactions were also found on +RBFox2 boundaries (Fig. 7W, Suppl. Fig. S9T-U, Suppl. Table S11). For example, decreased RBFox2 binding and intra-TAD interaction were found at the TAD boundary encompassing *Alyref2* gene (Fig. 7X). These results provide direct evidence to demonstrate that RBFox2 recruitment is facilitated by nascent transcription. In addition, we treated cells with RNase to remove the total RNAs or RNase inhibitor (RIN) to serve as the control (Suppl. Fig. S10A). A drastic reduction of RBFox2 binding number and landscape was observed on a global scale (Suppl. Fig. S10B-C). Most of the +RBFox2 boundaries were lost (Suppl. Fig. S10D) while over 70% of all boundaries remained stable upon RNase treatment (Suppl. Fig. S10E). Also, no significant impact on TAD insulation and genome organization was detected (Suppl. Fig. S10F-L). Altogether, the above findings from C2C12 support our conclusion that co-transcriptional binding of RBFox2 with nascent transcripts may have a role in TAD boundary formation and maintenance (Fig. 7Y).

## Discussion

In this study, by harnessing the available global DNA-binding profiles of RBPs from K562 and HepG2 cells we delineated the roles of dozens of RBPs in 3D genome organization in particular how they may participate in TAD organization. We demonstrate that TAD boundaries are hotspots for RBP binding and the enrichment is associated with increased TAD insulation strength and in a CTCF independent manner, implying a role for RBP in TAD organization (Fig. 6O). Using RBFox2 as an example for further investigation, we show that RBFox2 binding at TAD boundaries can indeed promote the insulation ability. And similar findings can extrapolate to C2C12 myoblast cells where we found RBFox2 regulation of TAD boundaries may be relevant in the process of myoblasts differentiation into myotubes (Fig. 7Y). Altogether, our findings have unveiled the pervasiveness of RBP involvement in 3D genome regulation and strengthened the emerging importance of RBPs in transcriptional regulation.

Although it is commonly accepted that cohesin-mediated extrusion preferentially restricted by CTCF-enriched boundaries are likely critical regulatory elements of TAD structure and boundary insulation, new possible regulatory mechanisms underlying TAD structure and function have become a focus of intense investigation within the field of 3D genome regulation. Findings from our study provide a previously unknown mechanism that is mediated by RBPs. We found that all the examined 33 RBPs in K562 and 23 in HepG2 display certain degree of enrichment on TAD boundaries and a large portion of all TAD boundaries contain at least one RBP. The prevalent association of RBPs with TAD boundaries indicates a potential role of RBPs in regulating TAD organization. A positive association of the binding with TAD insulation strength was found for each of the identified baRBPs. It is also likely that these baRBPs may function in a combinatorial mode as suggested by our network analyses, and there may also be synergistic interactions with TFs binding to the boundaries. Nevertheless, some baRBPs such as RBFox2 may play a more influential role than others. The TAD-regulating roles of baRBPs appear to be conserved in both K562 and HepG2 and can also be observed in C2C12, highlighting this as a universal phenomenon in cells. Considering the large repertoire of both conventional RBPs and the expanding list of novel types of unconventional RBPs [45, 46], we believe the number of baRBPs is much larger than what was identified using the limited RBP ChIP-seq datasets in our study, and their binding and functions may be cell context specific and dynamically regulated by various signaling [47].

Our findings also add new evidence for the emerging role of RBPs in transcription regulation. Several recent studies have unveiled a widespread RBP occupancy on open chromatin [23, 25, 31, 47] and attracted ongoing efforts in illuminating the functional meaning of the RBP-chromatin interaction. Despite the field remains nascent, mounting evidence supports that chrRBPs may have fundamental roles in transcriptional regulation via diverse mechanisms. Expectedly, some of our baRBPs overlap with the list of chrRBPs, suggesting they may bind to chromatin directly. It is interesting to identify the GC-rich sequences and possibly dG4s as binding elements for all the baRBPs. dG4s are a non-classical type of DNA secondary structures pervasively observed on gene promoters and it is believed that dG4s can act as recruiting platforms for TFs; emerging evidence also suggests the global enrichment of dG4s on TAD boundaries and their potential participation in TAD organization [48–50], it will thus be interesting to investigate whether baRBPs can be directly recruited to dG4s on the TAD boundaries or indirectly through TFs or other factors. It is also noteworthy to point out that certain chrRBP such as DDX41 directly recognizes R-loops which are unique RNA-DNA hybrid structures known to stabilize TAD boundaries [51].

Nevertheless, we believe that the baRBPs mainly depend on nascent promoter transcripts or active transcription to facilitate their binding and function on the boundaries. It is long accepted that despite CTCF and cohesin play a chief role in TAD formation in mammalian cells, a subset of TAD boundaries may be set by transcriptionally active genes in a CTCF independent manner [16]. It was speculated that highly transcribed genes might form a barrier to prevent loop extrusion by cohesin via accumulation of large amounts of transcriptional machinery and regulatory factors [52]. Here our findings provide additional insights by demonstrating that baRBPs may be active components of non-CTCF driven boundaries. Consistently, the enhancing effect of baRBPs was prominent on CTCF absent boundaries but not seen on CTCF present boundaries (Fig. 4); it is likely that baRBPs may be sufficient to block cohesin passively or even via direct physical interaction [53]. Alternatively, emerging evidence suggests that nuclear condensates can organize long-distance DNA-DNA interactions through surface tension [12] therefore phase separation could fulfill an important role in TAD organization. In this scenario, nascent transcripts on the TAD boundaries could act as a scaffold to recruit baRBPs that will generate condensates; to this end, we did predict the baRBPs are highly capable of forming condensates (data now shown). The RNA transcripts could also nucleate or facilitate condensates that subsequently enhance interactions between their respective TAD boundaries and intra-TAD loci. It is not clear whether the phase separation mediated boundary regulation requires cohesin and CTCF at all; it is likely that the two modes exist separately or synergistically depending on the composition and variances of RNA-baRBP complexes or scaffolds at TAD boundaries, which will be critical points of inquiry in understanding the mechanisms in more depth. It is also interestingly to note that TADs with +baRBP boundaries were significantly smaller in size, coincidently, in *Drosophila* embryos the TAD-like domains with active promoters on their boundaries are also small in size [17], indicating the diversified nature of TADs depending on the organizing force/factors.

In fact, it is well known that noncoding RNAs accumulated on TADs play important structural and regulatory roles [12, 54], our studies detect that promoter mRNA transcripts could be the primary force for baRBP interaction and TAD organization. Of note, a recent study suggests that nascent transcripts can facilitate the phase separation of RBPs, which confines Pol II to the proximity of transcription sites to promote transcription [25]. Therefore, in theory various forms of nascent RNAs could be active participants of gene regulation by harnessing their convenient proximity to the chromatin and co-transcriptional enlisting capacity of baRBPs. In the future, it will be interesting to further investigate the potential interaction of other chromatin associated RNA species such as eRNAs [27, 55] or retrotransponson derived transcripts [56] with baRBPs on TAD boundaries. Together with nascent transcripts, baRBPs could execute transcriptional regulatory functions through diversified means such as via impacting TAD organization or modulating Pol II activities. Our findings thus highlight the emerging unconventional functions of RBPs beyond their traditional framework in regulating RNA metabolism and unveil the intimate interplay between transcriptional and co-transcriptional regulations.

After uncovering the positive association of baRBP binding with TAD insulation ability, it is imperative to test the causative relationship. We thus targeted RBFox2 for further functional and mechanistic elucidation. RBFox2 is one of the few RBPs with well documented function in transcriptional regulation. The study from Wei C et. al. [57] provides solid evidence that RBFox2 has a direct role in transcriptional control besides splicing regulation; nascent RNAs are required for RBFox2 interaction with chromatin where it directly interacts with PRC2 to mediate dynamic transcriptional control. Our findings thus expand the mode through which RBFox2 can regulate transcription. When depleting *RBFox2* from K562 cells, however, no marked impact on TAD insulation ability was observed despite altered intra-and inter-TAD interactions; and in C2C12 cells its depletion surprisingly caused enhancing effect on TAD insulation. We therefore suspect RBFox2 alone may not be sufficient and it requires the cooperative actions of many baRBPs and factors for example POLR2G, GTF2F1 and HNRNPL (Fig. 2). Our results from C2C12 cells also demonstrate that RBFox2 regulation of TAD organization is relevant in the process of MB differentiation to MT, pointing to a potential role of RBFox2 in skeletal muscle development and regeneration which can be further investigated in the future. Additionally, our findings (Fig. 5) also confirmed the corporation of RBFox2 with nascent transcription. Interestingly, transcriptional inhibition by ActD treatment drastically attenuated RBFox2 binding and TAD insulation ability, while RNA removal by RNase A treatment decreased RBFox2 binding but had no obvious effect on boundary insulation, indicating that it is likely the nascent transcription not the RNAs are required. Alternatively, RNase A treatment, which targets only free ssRNAs, might not immediately inflict a loss of TAD insulation which is moderately resilient to the loss. We predict that the sophisticated regulation of TAD formation/organization by baRBPs, transcripts and other factors will be the focus of intense investigation in the coming years, which will shed novel lights on the gene regulatory mechanisms in cells.

## Methods

### Cells and transfection

Mouse C2C12 myoblast cells (CRL-1772) were obtained from American Type Culture Collection (ATCC) and cultured as described before [24, 58] in DMEM medium (Gibco, 12800-017) with 10% fetal bovine serum (Gibco, 10270-106), 100 units/ml of penicillin, and 100 μg of streptomycin at 37°C in 5% CO_2_. Human leukemia K562 cells (CCL-243) were obtained from ATCC and maintained at 37°C under 5% CO_2_ in RPMI-1640 medium (Thermo Fisher Scientific, 22400089) with 10% fetal bovine serum, 100 units/ml of penicillin, and 100μg of streptomycin. All cell lines were tested as negative for mycoplasma contamination. For transfection, siRNA against *RBFox2* or control oligos were obtained from Shanghai GenePharma Corp., China. Transfections of K562 cells were conducted with Amaxa Cell Line Nucleofector Kit V (Lonza, Basel, Switzerland). Briefly, K562 cells were grown for 24h and resuspended in 100μl Nucleofector Solution V. siRNA oligos targeting *RBFox2* or control oligos were used at a concentration of 300nM and the transfection was completed with Lonza Nucleofector^™^ using program T-016. Sequences of siRNA oligos are included in Suppl. Table S12.

### CRISPR/Cas9 mediated knock-out (KO)

For generating C2C12 cells with *Rbfox2* KO, two site-specific single guide RNAs (sgRNAs) targeting the exon 2 of *Rbfox2* isoform 6 (NM_001110830.3 from National Center for Biotechnology Information) in mice were selected using a web tool Crispor (http://crispor.tefor.net/) and cloned into pX330-EGFP plasmid respectively (42230, Addgene). As described before [59], C2C12 cells were simultaneously transfected with the two plasmids by Lipofectamine 3000 and Flow cytometry was employed for sorting out EGFP+ cells after 48h of transfection; the EGFP+ cells were then diluted in 96-well plates to obtain single-cell clones. Each colony was subject to PCR genotyping until the KO clones were identified. Sequences of the primers for genotyping and sgRNAs are listed in Suppl. Table S12.

### Actinomycin-D and RNase A treatment

For transcriptional inhibition, around 1 × 10^7^ C2C12 myoblasts were treated with actinomycin D (1 μg/ml) while culturing for 24 h then harvested for Hi-C crosslinking. For RNA degradation, 1 × 10^7^ C2C12 myoblasts were subject to RNase A or RNase inhibitor (RIN) treatment. Cells were spun down at 2,000 × g for 10 min at 4°C, t and the supernatant was removed. The cell pellet was re-suspended with 1 ml of cold PBS + 0.02% Tween-20 followed by incubation on ice for 10 min. Next, either 35U of RNase A (Invitrogen, 12091-021) or 30U of RIN (Life Technologies, 10777019) was added to the suspension and incubated at 37°C for 30 min in a thermomixer at 950 rpm. After incubation, the cells were put on ice for 10 min then spun down at 2,000 × g for 10 min at 4°C and washed once with 1 ml cold PBS with protease inhibitor cocktail (Cell Signaling Technology, 7012L).

### RNA isolation and RT-PCR

As described before [60, 61], total RNAs were extracted using TRIzol reagent (Invitrogen) according to the manufacturer’s instruction. For RT-PCR, cDNAs were synthesized under the manufacturer’s instructions of HiScript® II Reverse Transcriptase Kit with gDNA wiper (Vazyme, R223-01). Real-time PCR was performed to quantify the expression level of mRNAs by Luna® Universal qPCR Master Mix (NEB, M3003E) and LightCycler ®480 Real-Time PCR System (Roche). GAPDH (glyceraldehydes 3-phosphate dehydrogenase) or 18s RNA were used as internal controls for normalization. Sequences of all primers used are listed in Suppl. Table S12. The relative fold changes compared to control groups were calculated by the classical ΔΔCt method.

### Western blotting

As described before [62, 63], cell lysates were collected by direct lysis with RIPA buffer (50 mM Tris-HCl, PH 7.5, 150 mM NaCl, 1.0 mM EDTA, 0.1% SDS, 1% Sodium deoxycholate and 1% Triton X-100) supplemented with protease inhibitor cocktail (PIC, 88266, Thermo Fisher Scientific), samples were boiled to denature at 100°C for 10 min before SDS-PAGE gel electrophoresis. The proteins were wet-transferred to PVDF Western blotting membrane (03010040001, Roche) after electrophoresis, membranes were blocked for 1 h with 5% milk in TBST before overnight incubation with below primary antibodies:RBFox2 (Bethyl Laboratories, A300-864A), GAPDH (ABS16, Sigma-Aldrich). Membranes were then washed three times and incubated with Goat anti-rabbit IgG-HRP, Goat Secondary Antibody (ABclonal, AS014) or Anti-goat anti-mouse IgG-HRP, Mouse Secondary Antibody (ABclonal, AS003) for 1 h at RT. Reactive proteins were detected by Enhanced Chemiluminescence (ECL) reagent (K-12045-D20, Advansta).

### ChIP-seq

The RBFox2 ChIP-seq libraries in C2C12 cells were prepared as previously reported [25]. Briefly, both formaldehyde and DSP (dithiobis succinimidyl propionate) were used to dual cross-link the proteins to the DNA. Approximately 1 × 10^7^ C2C12 cells were cross-linked with 0.2 mM DSP for 30 min at room temperature, combined with a 15-minute 3% final concentration formaldehyde treatment. Cells were then quenched with 1/10 volume of 2.5 M Glycine and washed and collected by centrifugation at 2,000 × g for 5 min at 4°C, flash-frozen in liquid nitrogen, and stored at-80°C. Next, following our standard procedure, 6 μg of antibody against RBFox2 (Bethyl Laboratories, A300-864A) was used for immunoprecipitation. Immunoprecipitated genomic DNA was resuspended in 25 μl of water. For DNA library construction, a NEBNext® Ultra™ II DNA Library Prep Kit for Illumina® (NEB, E7645S) was used according to the manufacturer’s instructions. Bioanalyzer analysis and qPCR were used to measure the quality of DNA libraries including the DNA size and purity. Finally, DNA libraries were sequenced on the Illumina Genome Analyzer II platform. [64]

### In situ Hi-C

The in situ Hi-C libraries were prepared as previously reported [42]. About 300,000 C2C12 cells were cross-linked and processed to generate Hi-C libraries. The samples were lysed in 500 μl of ice-cold Hi-C lysis buffer [10 mM tris-HCl (pH 8.0), 10 mM NaCl, and 0.2% Igepal CA630 supplemented with complete proteinase inhibitors] on ice for 15 min. Samples were then centrifuged at 2500g for 5 min. Pelleted nuclei were washed once with 500 μl of 1.25× NEBuffer 3.1. The supernatant was discarded, the nuclei were resuspended in 358 μl of 1.25× NEBuffer 3.1, and 11 μl of 10% SDS was added followed by incubation at 37°C for 1 hour. After incubation, 75 μl of 10% Triton X-100 was added to quench the SDS and then incubated at 37°C for 1 hour. One hundred units of Dpn II restriction enzyme (NEB, R0543) was added, and chromatins were digested at 37°C for overnight. Samples were incubated at 62°C for 20 min to inactivate Dpn II and then cooled to room temperature. Samples were then centrifuged at 2500g for 5 min. Pelleted nuclei were resuspended in 50 μl of fill-in master mix [3.75 μl of 0.4 mM biotin-14-dATP (2′-deoxyadenosine-5′-triphosphate), 1.5 μl of 1 mM 2′-deoxycytidine-5′-triphosphate (dCTP), 1.5 μl of 1 mM 2′-deoxyguanosine-5′-triphosphate (dGTP), 1.5 μl of 1 mM 3′-deoxythymidine-5′-triphosphate (dTTP) mix, 2 μl of DNA polymerase I (5 U μl^−1^), Large (Klenow) Fragment, and 1× NEBuffer 3.1] to fill in the restriction fragment overhangs and mark the DNA ends with biotin. Samples were mixed by pipetting and incubated at 23°C for 4 hours. Ligation master mix [398 μl of water, 50 μl of 10× NEB T4 DNA ligase buffer, 1 μl of bovine serum albumin (50 mg ml^−1^), 1 μl of T4 DNA ligase (400 U μl^−1^)] was added, and samples were incubated at 16°C for overnight in ThermoMixer C with interval shake. Nuclei were pelleted by centrifugation at 2500g for 5 min, and 380 μl of the supernatant was discarded. Pellets were then resuspended in the remaining 120 μl of ligation mix supplemented with 12 μl of 10% SDS and 5 μl of proteinase K (20 mg ml^−1^) and incubated at 55°C for 2 hours, with shaking at 1000 rpm. Fifteen microliters of 5 M NaCl was added, and the reaction was incubated at 65°C for 16 hours. DNA samples were subjected to phenol:chloroform extraction and ethanol precipitation and finally resuspended in 130 μl of 0.1× TE buffer. The purified DNA samples were then sheared to a length of ∼300 bp using Covaris S220 instrument (intensity:175 W, duty cycle:10%, cycles per burst:200, time:150 s). The biotinylated DNA was pulled down by 10 μl of Dynabeads MyOne Streptavidin C1 beads (10 mg ml^−1^; Life Technologies, 65001). The beads were then resuspended in 23 μl of 10 mM tris-Cl (pH 8.0), and libraries were prepared by on-bead reactions using the NEBNext Ultra II DNA Library Preparation Kit (NEB, E7645S). The beads were separated on a magnetic stand, and the supernatant was discarded. After washes, the beads were resuspended in 20 μl of 10 mM tris buffer and boiled at 98°C for 10 min. The elute was amplified for 10 to 13 cycles of PCR with Phanta Master Mix (Vazyme, P511-01), and the PCR products were purified using VAHTS DNA Clean Beads (Vazyme, N411-01). Libraries underwent paired-end sequencing on an Illumina HiSeq X Ten instrument.

### Gene expression analysis of RNA-seq data

The raw reads of total RNA-seq were processed following the procedures described in our previous publication [61, 65–67]. Briefly, the adapter and low-quality sequences were trimmed from 3’ to 5’ ends for each read and the reads shorter than 50 bp were discarded. The reads that passed the quality control were mapped to human (hg19) or mouse genome (mm9) with Bowtie2 [68]. Cufflinks was then used to estimate transcript abundance in Fragments Per Kilobase per Million (FPKM) [69]. Genes were annotated as differentially expressed if the change of expression level is greater than 2 folds between two stages/conditions.

### ChIP-seq data analysis

For analyzing the RBFox2 ChIP-seq data, raw reads were processed as previously described [70] Briefly, the adapter and low-quality sequences were trimmed from 3’ to 5’ ends by Trimmomatic [71] and the reads shorter than 36 bp were discarded. Subsequently, the preprocessed reads were aligned to the mouse genome (mm9) using Bowtie2. The aligned reads were then converted to bam format using samtools, and the duplicate reads were removed by Picard (http://broadinstitute.github.io/picard). Peaks were then identified by MACS2 with q-value equal to 0.01 by using the IgG control sample as background. For the published RBP ChIP-seq data, motif enrichment analysis was performed on the ChIP-seq peaks residing at TAD boundaries by using software DREME with default parameters [72].

### NMF Analysis

A unifying set of cis-acting regulatory elements (CREs) was defined based on ChIP-seq peaks for RBPs and TFs. ChIP-seq peaks with the distance from their summits ≤ 1 Kb were merged into one CRE by Bedtools. A matrix was then built, with 1/0 representing whether a CRE (row) was occupied or not by each TF/RBP (column). CREs occupied by multiple factors were then subject to NMF analysis as described [69]. The rank of 5 and 4 was selected based on both cophenetic correlation coefficient (CPCC) and dispersion coefficient in K562 and HepG2 cells, respectively. NMF was then run with the selected rank. Protein-protein physical interactions were obtained from GeneMANIA (http://genemania.org).

### In situ Hi-C data processing

Raw Hi-C data were processed as previously described [42, 59]. Briefly, the in-situ Hi-C data was processed with a standard pipeline HiC-Pro [29]. First, adaptor sequences and poor-quality reads were removed using Trimmomatic (ILLUMINACLIP:TruSeq3-PE-2.fa:2:30:10; SLIDINGWINDOW:4:15; MINLEN:50). The filtered reads were then aligned to human (hg19) or mouse reference genome (mm9) in two steps:1) global alignment was first conducted for all pair-end reads, 2) the unaligned reads were split into prospective fragments using restriction enzyme recognition sequence (GATCGATC) and aligned again. All aligned reads were then merged and assigned to restriction fragment, while low quality (MAPQ<30) or multiple alignment reads were discarded. Invalid fragments including unpaired fragments (singleton), juxtaposed fragments (re-ligation pairs), un-ligated fragments (dangling end), self-circularized fragments (self-cycle), and PCR duplicates were removed from each biological replicate. The remaining validate pairs from all replicates of each stage were then merged, followed by read depth normalization using HOMER (http://homer.ucsd.edu/homer/ngs/) and matrix balancing using iterative correction and eigenvector decomposition (ICE) normalization to obtain comparable interaction matrix between different stages.

### Identification and analysis of compartments and TADs

Following previous procedure [10], to separate the genome into A and B compartments, the ICE normalized intra-chromosomal interaction matrices at 100-kb resolution were transformed to observe/expect contact matrices, and the background (expected) contact matrices were generated to eliminate bias caused by distance-dependent decay of interaction frequency and read depth difference[42, 59]. Pearson correlation was then applied to the transformed matrices and the first principal component (PC1) of these matrices was divided into two clusters. The annotation of genes and the expression profile were used to assign positive PC1 value to gene-rich component as compartment A and negative PC1 value to gene-poor component as compartment B. The compartmentalization strength was measured using cooltools (https://github.com/mirnylab/cooltools). Briefly, all 100-kb compartment bins were divided into 50 equal degrees based on the ranking of PC1 values, and the average interaction strength (observed/expected) was calculated between pairs of 100-kb loci arranged by their PC1 and normalized by genomic distance. The dynamics of interaction strength for intra/inter compartment (A vs. A, B vs. B, A vs. B) were measured using top 20% of compartment bins from both compartments A and B based on absolute PC1 value. Normalized contact matrix at 10 kb resolution of each time point was used for TAD identification using TopDom [30]. In brief, for each 10-kb bin across the genome, a signal of the average interaction frequency of all pairs of genome regions within a distinct window centered on this bin was calculated, thus TAD boundary was identified with local minimal signal within certain window. The false detected TADs without local interaction aggregation were filtered out by statistical testing. Invariant TADs were defined using following criteria:1) the distance of both TAD boundaries between two conditions is no more than 10 kb; 2) the overlapping between two TADs should be larger than 80%; stage-specific TADs were defined otherwise. The insulation score of the identified TAD boundary was also defined as previously described [73], which used the local maximum on the outside of TAD to minus the local minimum on the inside of TAD of each boundary bin. The domain score was calculated for each TAD at 10 kb interaction matrix through dividing total intra-TAD contacts by all contacts involving the TAD.

### Aggregate analysis

ADA and APA were performed by using coolpuppy [74]. Aggregate analysis of interaction frequency around boundaries were processed by AnalyzeHiC module of HOMER and then visualized by Java Treeview (https://jtreeview.sourceforge.net/).

### GRO-seq analysis

The processing of GRO-seq is described in previous paper [24]. Briefly, low-quality reads were filtered out and adaptor sequences were trimmed from raw reads using Trimmomatic-0.3658. The remaining reads were then aligned to the mouse genome (mm9) using Bowtie2. If multiple reads aligned to the same genomic position, only one read per position was kept for downstream analyses. Primary transcripts were de novo identified throughout the genome using HOMER. To define putative eRNAs, transcripts overlapping with protein-coding genes, antisense transcripts, divergent transcripts and the other genic regions (rRNA, snRNA, miRNA, snoRNA, etc.) were filtered and the remaining transcripts were defined as putative eRNAs if their de novo identified transcriptional start site (TSS) was located in super-enhancer or typical-enhancer regions. The quantification of gene expression was processed by analyzerepeat module of HOMER.

### Meta-gene analysis

The distribution of ChIP-seq or GRO-seq signal around regions of interest were performed by using ngsplot software [75] with parameters-N 0.5-XYL 0-LEG 0-BOX 0-VLN 0.

### Data availability

In situ Hi-C and ChIP-seq data reported in this paper are deposited in the Gene Expression Omnibus database under accession GSE243534. All used datasets from other publications are summarized in Suppl. Table S1. All other data supporting the findings of this study are available from the corresponding author on reasonable request.

## Acknowledgments

We thank Dr. Yafei Yin (Zhejiang University, China) for providing RBP ChIP-seq protocol. We thank Dr. Xiaohua Shen (Tsinghua University, China) for her helpful comments and suggestions. This work was supported by National Key R&D Program of China to H.W. (project code:2022YFA0806003); General Research Fund (GRF) from Research Grants Council (RGC) of the HongKong (HK) Special Administrative Region, China to H.W. (project codes:14100620, 14105823, 14106521 and 14115319 to H.W.; 14105123, 14103522, 14120420 and 14120619 to H.S.); Theme-based Research Scheme (TRS) from RGC (project code:T13-602/21-N); Collaborative Research Fund (CRF) from RGC (project code:C6018-19GF); Health and Medical Research Fund (HMRF) from Health Bureau of HK to H.W. (project codes:10210906 and 08190626); the National Natural Science Foundation of China (NSFC) to H.W. (project codes:82172436 and 31871304); the research funds from Health@InnoHK program launched by Innovation Technology Commission, the Government of HK to H.W.; CUHK Strategic Seed Funding for Collaborative Research Scheme (SSFCRS) to H.W.; Area of Excellence Scheme (AoE) from RGC (project code:AoE/M-402/20).

## Author Contributions

H.W., Q.S. and H.S. designed the project; Q.Z., and Y.Q. generated library of high-throughput sequencing data; Q.S. performed bioinformatic analyses; Q.S. and H.W. wrote the paper. All authors read and approved the final manuscript.

## Inventory of Supplementary Information

### 1. Supplementary Figures

Figure S1. TAD boundaries are hotspots for RBP binding.

Figure S2. Network interaction of baRBPs and TFs at TAD boundaries.

Figure S3. Comparison of insulation scores of +baRBP boundaries and-baRBP boundaries.

Figure S4. baRBP enrichment correlates with increased insulation strength of TAD boundaries.

Figure S5. Comparison of insulation scores of +baRBP boundaries and-baRBP boundaries at CTCF present or absent boundaries.

Figure S6. baRBPs may facilitate TAD organization independent of CTCF and cohesin binding.

Figure S7. baRBP enrichment at TAD boundaries is correlated with active transcription.

Figure S8. RBFox2 binding promotes TAD organization in K562.

Figure S9. RBFox2 regulation of 3D genome is relevant in mouse myoblast differentiation.

Figure S10. RNase treatment have moderate effect on 3D genome organization.

### 2. Supplementary Tables

Supplementary Table S1. Sources of high throughput sequencing data used in the study.

Supplementary Table S2. Lists of TADs and TAD boundaries identified in K562 and HepG2 cells.

Supplementary Table S3. Network analysis of baRBPs on TAD boundaries.

Supplementary Table S4. Analysis of TADs with baRBP binding boundaries.

Supplementary Table S5. Analysis of RBP co-binding with CTCF and cohesin.

Supplementary Table S6. Analysis of GRO-seq signals at baRBP enriched boundaries.

Supplementary Table S7. baRBP ChIP-seq peaks at promoter containing boundaries.

Supplementary Table S8. Analysis of RBFox2 knock-down effect on 3D genome organization in K562 cells.

Supplementary Table S9. Analysis of TAD and TAD boundary in C2C12 MB and MT cells.

Supplementary Table S10. Analysis of RBFox2 knock-out effect on 3D genome organization in C2C12 cells.

Supplementary Table S11. Analysis of the effect of ActD or RNase treatment on 3D genome.

Supplementary Table S12. Sequences of oligos used in this study.

### 3. Supplementary Figures Legends

**Supplementary Figure S1.**
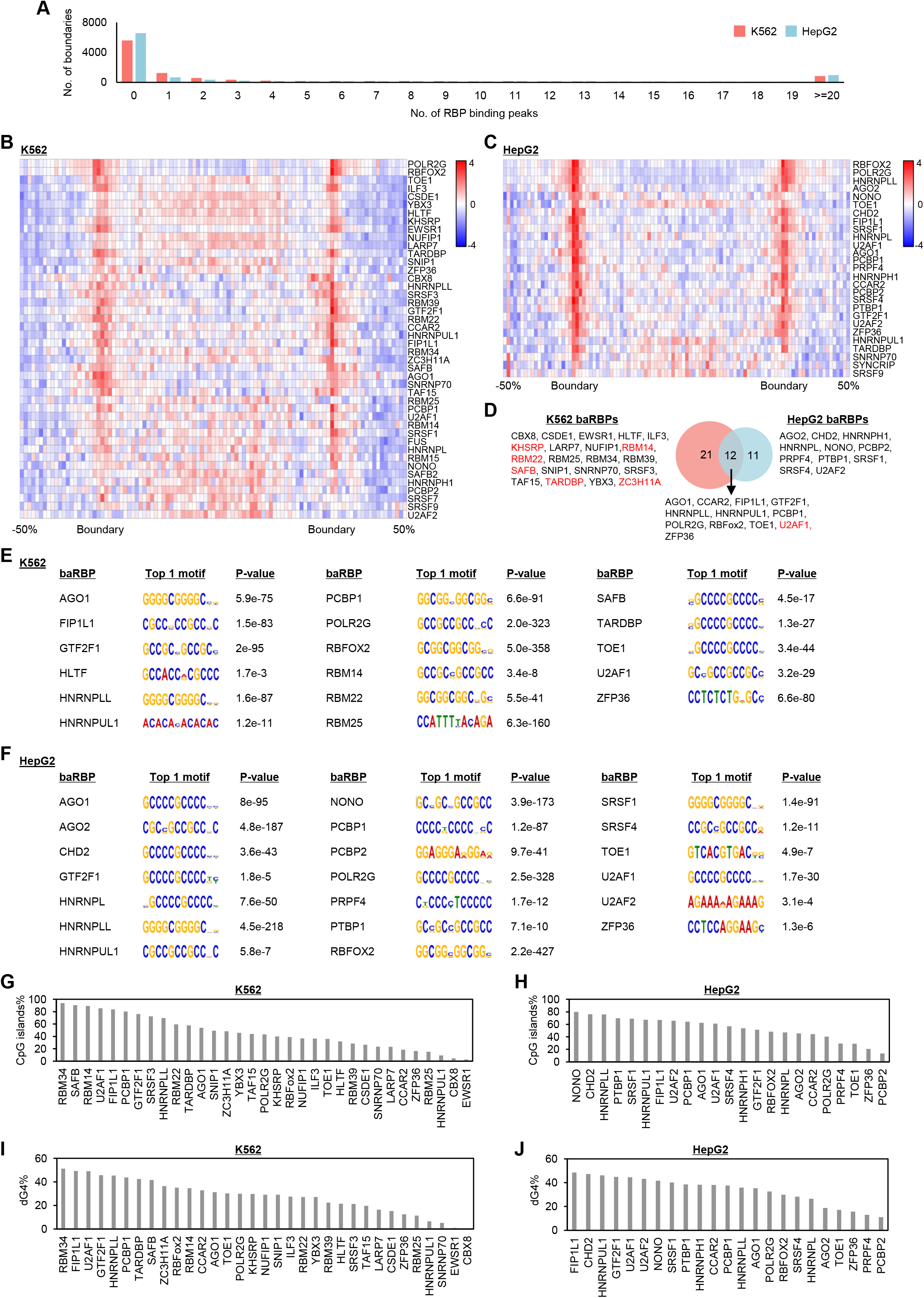
TAD boundaries are hotspots for RBP binding. **(A)** The number of boundaries with the indicated number of RBP binding peaks. **(B-C)** Heatmaps showing the RBP ChIP-seq signals around TADs in K562 and HepG2 cells. The ±50% region flanking the TADs was used for profiling. **(D)** Overlapping of baRBPs in K562 and HepG2 cells. Known chromatin associated RBP (chrRBPs) are highlighted in red. **(E-F)** Enriched motifs of baRBP ChIP-seq peaks within TAD boundaries in K562 and HepG2 cells. **(G-H)** Bar plot showing the percentage of baRBP ChIP-seq peaks overlapped with CpG islands in K562 and HepG2 cells. **(I-J)** Bar plot showing the percentage of baRBP ChIP-seq peaks overlapped with dG4 ChIP-seq peaks in K562 and HepG2 cells.

**Supplementary Figure S2.**
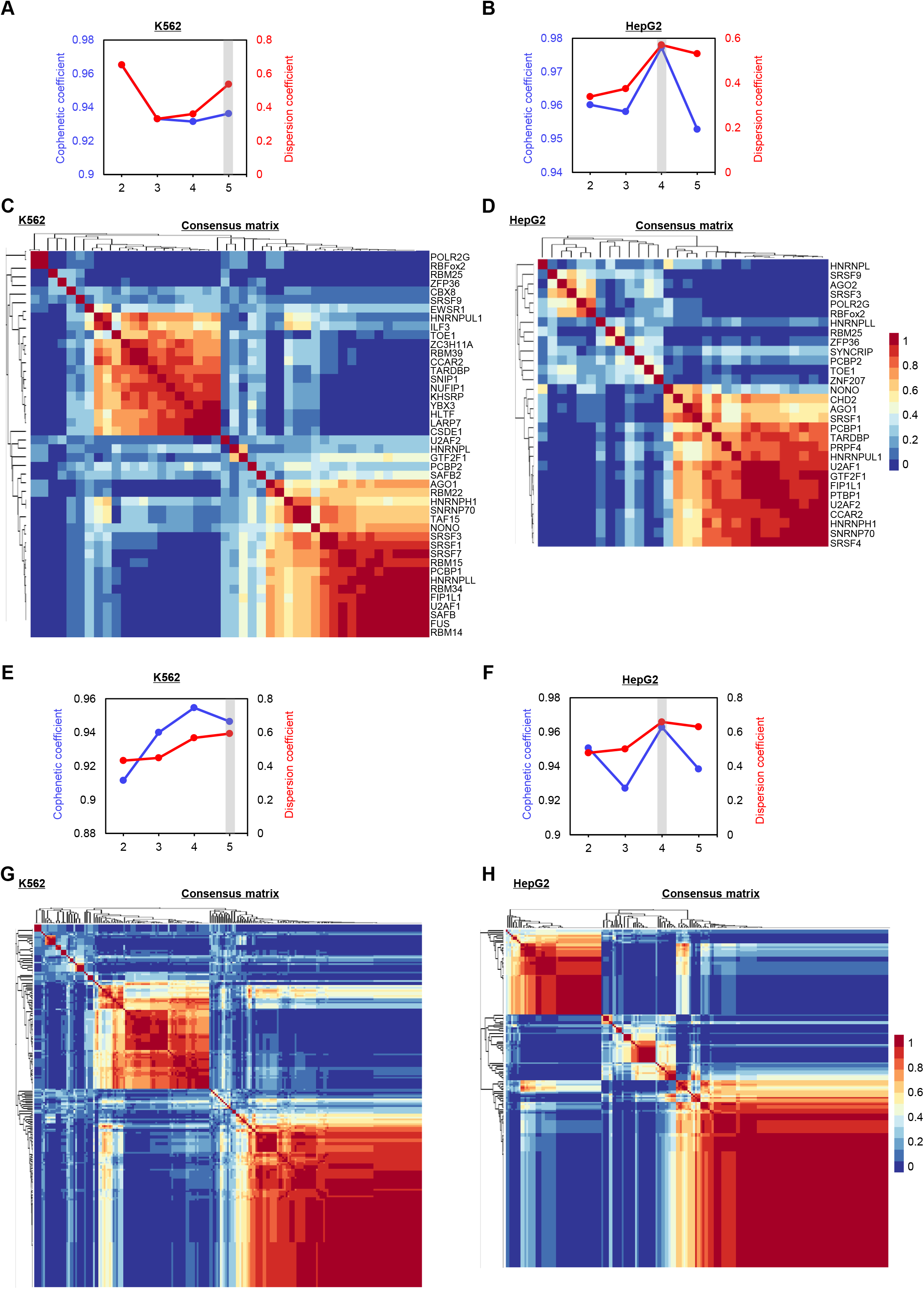
Network interaction of baRBPs and TFs at TAD boundaries. **(A-B)** Criteria for estimating a maximal stable factorization rank in NMF analysis for baRBP clusters in K562 and HepG2 cells. Grey line:the factorization rank of 5 and 4 was chosen for the representation of RBP groups in Figure 2B-C. **(C-D)** The average connectivity matrix in NMF analysis in Figure 2B-C, showing multiple groups that are distinct from each other. **(E-F)** Criteria for estimating a maximal stable factorization rank in NMF analysis for baRBP-TF clusters. Grey line:the factorization rank of 5 and 4 was chosen for the representation of baRBP groups in Figure 2F-G. **(G-H)** The average connectivity matrix in NMF analysis in Figure 2F-G, showing multiple groups that are distinct from each other.

**Supplementary Figure S3.**
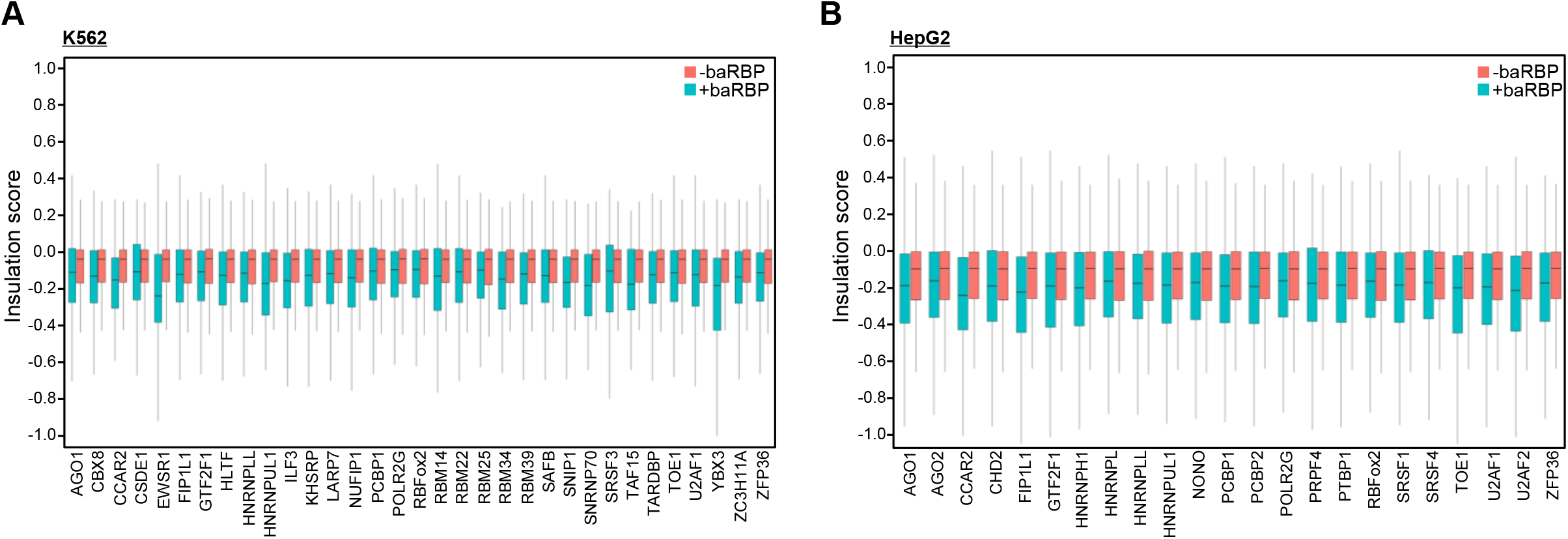
Comparison of insulation scores of +baRBP boundaries and-baRBP boundaries. **(A-B)** Comparison of insulation scores of boundaries with and without each of the baRBP (33 in K562 and 23 in HepG2).

**Supplementary Figure S4.**
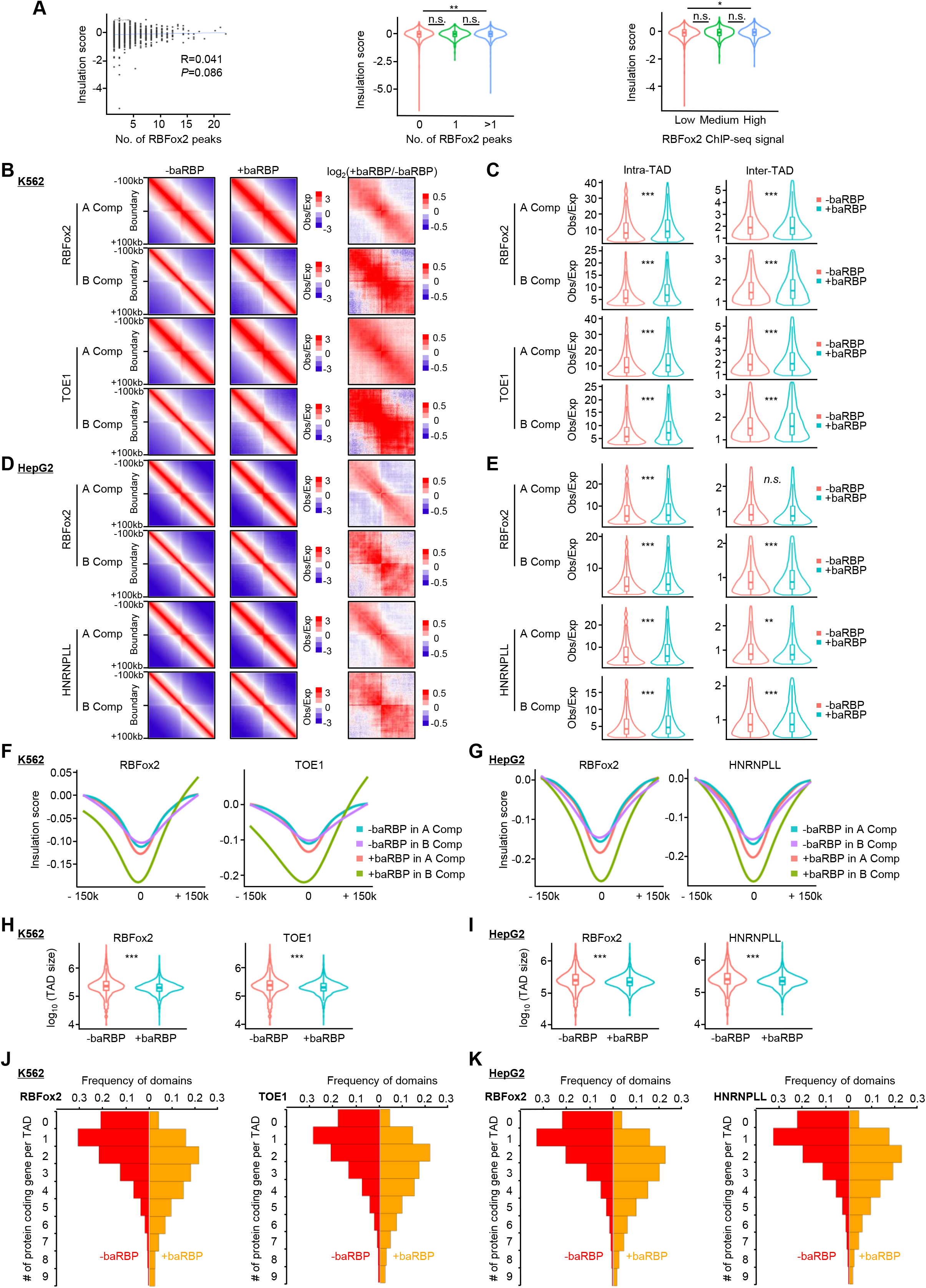
baRBP enrichment correlates with increased insulation strength of TAD boundaries. (**A)**) Correlation between insulation score and number of RBFox2 peaks (left and middle panels), and RBFox2 binding strength (low, medium or high) (right panel). **(B)** Aggregate analysis of interaction frequency around +baRBP (RBFox2 and TOE1) and-baRBP boundaries residing at A or B compartment in K562 cells. **(C)** Quantification of intra-or inter-TAD interaction frequency of data from **(B)**. **(D-E)** The above analyses were performed on baRBPs (RBFox2 and HNRNPLL) in HepG2 cells. **(F-G)** Comparison of insulation scores of the above +baRBP and-baRBP boundaries at A or B compartment in K562 and HepG2 cells. **(H-I)** Comparison of TAD size with or without the above +baRBP boundaries in K562 and HepG2 cells. **(J-K)** Comparison of numbers of genes per TAD with or without the above +baRBP boundaries in K562 and HepG2 cells.

**Supplementary Figure S5.**
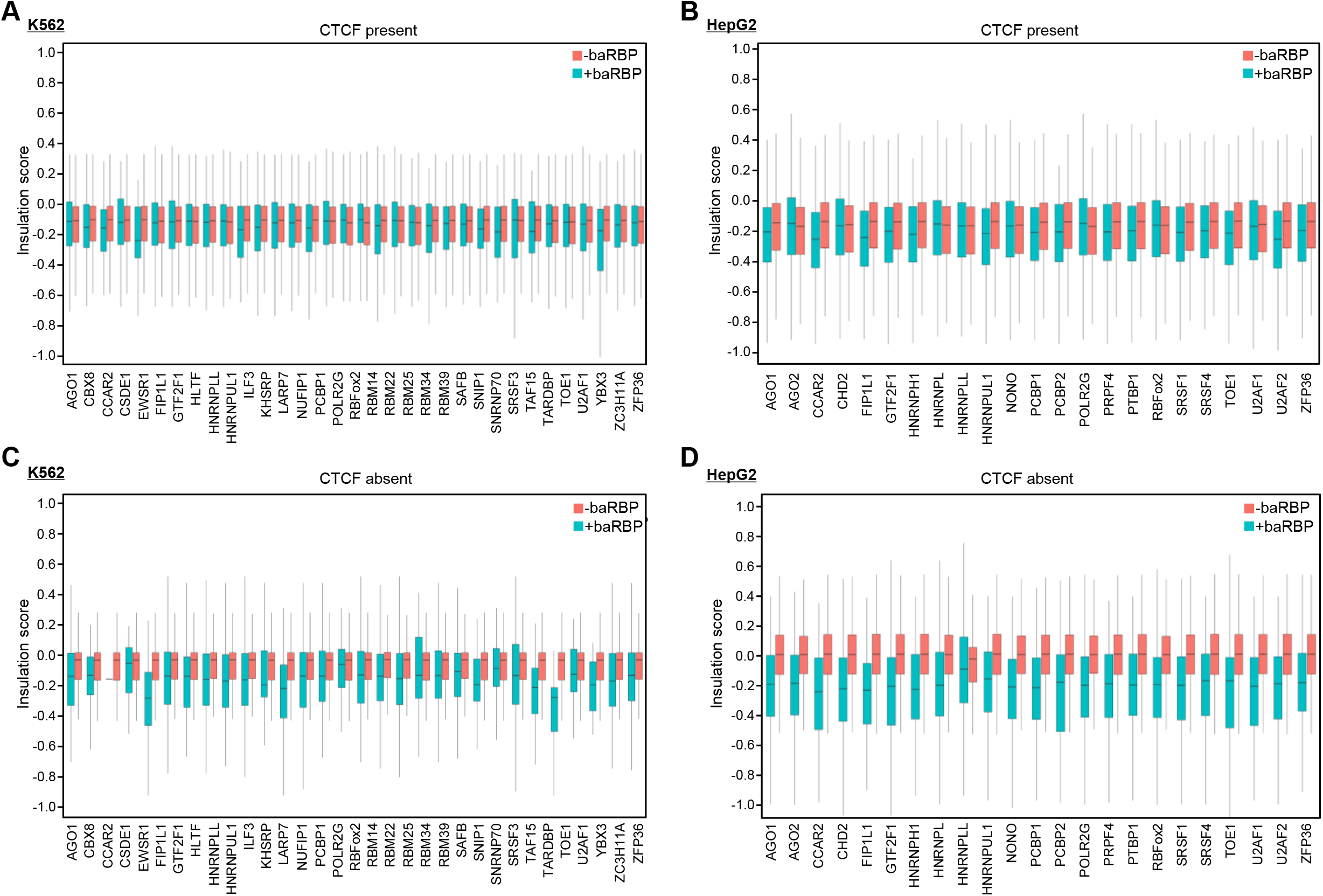
Comparison of insulation scores of +baRBP boundaries and-baRBP boundaries at CTCF present or absent boundaries. **(A-B)** Comparison of insulation scores of boundaries with and without each of the baRBP (33 in K562 and 23 in HepG2) binding at CTCF present boundaries. **(C-D)** Comparison of insulation scores of boundaries with and without the above baRBP binding at CTCF absent boundary in K562 and HepG2.

**Supplementary Figure S6.**
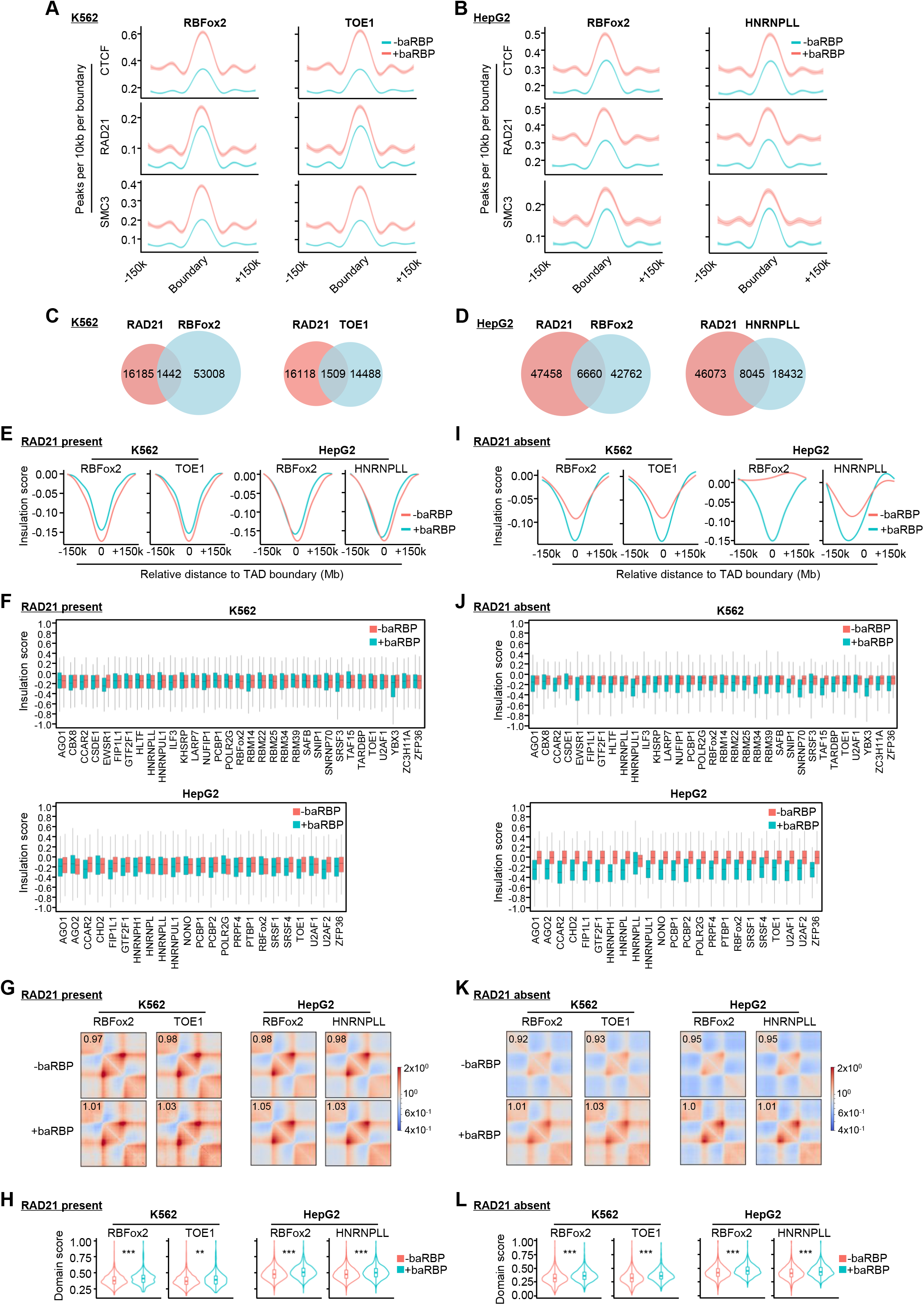
baRBPs may facilitate TAD organization independent of CTCF and cohesin binding. **(A-B)** CTCF, RAD21, and SMC3 peak counts around +baRBP (RBFox2 and TOE1 in K562, RBFox2 and HNRNPLL in HepG2) or-baRBP boundaries. **(C-D)** Overlapping of RAD21 and representative baRBP (RBFox2 and TOE1 in K562; RBFox2 and HNRNPLL in HepG2) ChIP-seq peaks at TAD boundaries. **(E)** Comparison of insulation scores of +baRBP (RBFox2 and TOE1 in K562 and RBFox2 and HNRNPLL in HepG2) and-baRBP boundaries at RAD21 present boundaries. **(F)** Comparison of the insulation scores of boundaries with and without each of baRBP (33 in K562 and 23 in HepG2). **(G)** Aggregate domain analysis of TADs with +baRBP (RBFox2 and TOE1 in K562 and RBFox2 and HNRNPLL in HepG2) and-baRBP boundaries at RAD21 present boundaries. **(H)** Comparison of domain scores of the above TADs with +baRBP and-baRBP boundaries at RAD21 present boundaries. **(I-L)** The above analyses were performed at RAD21 absent boundaries.

**Supplementary Figure S7.**
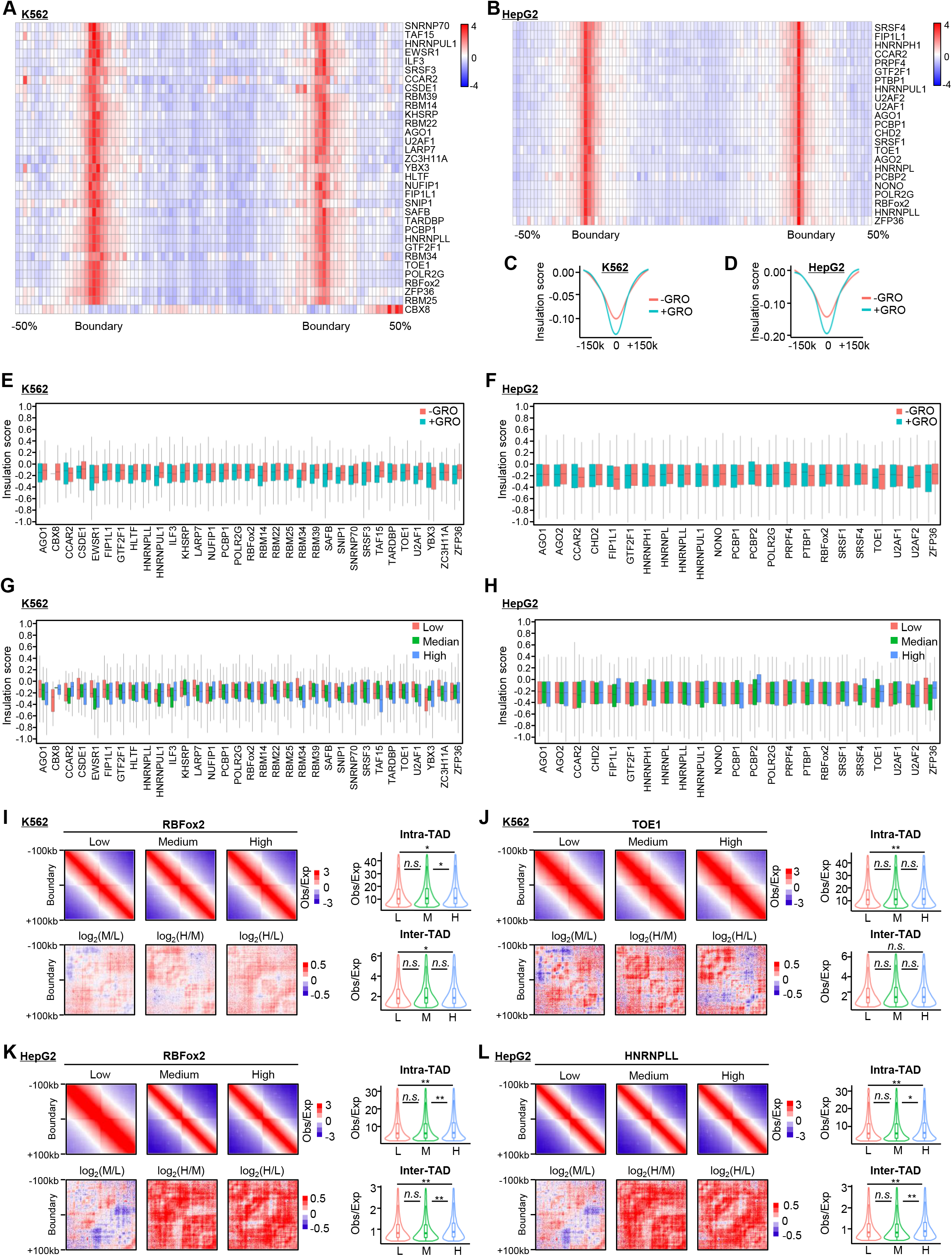
baRBP enrichment on TAD boundaries is correlated with active transcription. **(A-B)** Heatmaps showing the GRO-seq signals around TADs with +baRBP (33 in K562 and 23 in HepG2) boundaries. The ±50% region flanking each TAD was used for profiling. **(C-D)** Comparison of insulation scores of TAD boundaries with or without GRO-seq signals. **(E-F)** Comparison of insulation scores of each of +baRBP (33 in K562 and 23 in HepG2) boundaries with or without GRO-seq signals. **(G-H)** Comparison of insulation scores of the above +baRBP boundaries with low, medium, or high level of GRO-seq signals. **(I-J)** Aggregate analysis of interaction frequency around +baRBP boundaries (RBFox2 and TOE1) with low, medium or high level of GRO-seq signals in K562. **(K-L)** The above analyses were performed on RBFox2 and HNRNPLL in HepG2.

**Supplementary Figure S8.**
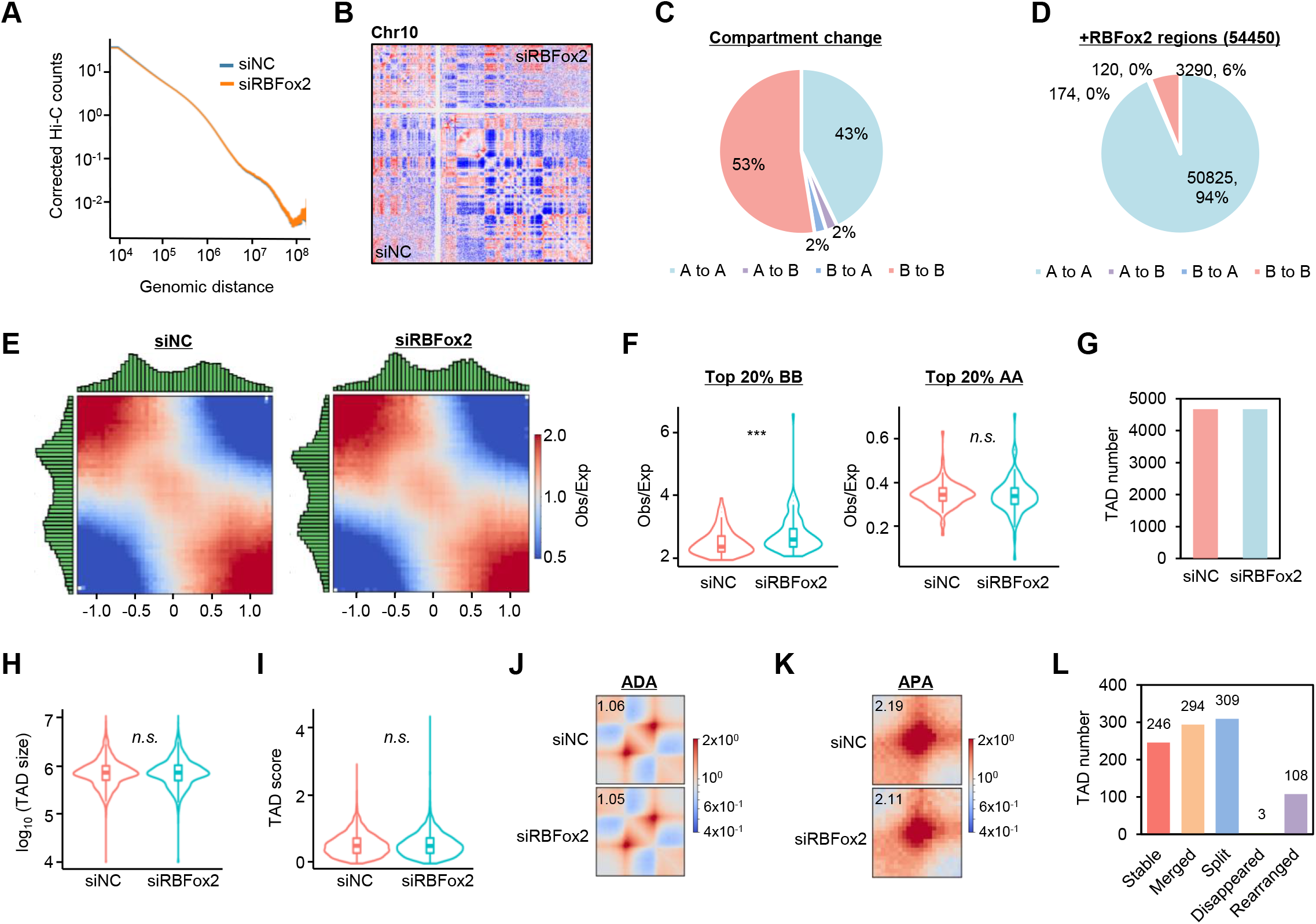
RBFox2 binding promotes TAD organization in K562. **(A)** Contact frequency as a function of genomic distance along the whole genome in *siNC* vs *siRBFox2* cells. **(B)** Representative heatmap showing the called compartment at chr10 in *siNC* (lower triangle) vs *siRBFox2* (upper triangle) cells. **(C)** Pie chart showing the fraction of genome with the designated type of compartment switching upon RBFox2 knock-down. **(D)** Pie chart showing the percentage of RBFox2 ChIP-seq peaks residing at each type of the above switched compartments. **(E)** Saddle plot showing the compartmentalization change in *siNC* vs *siRBFox2* cells. **(F)** Comparison of top 20% A-A or B-B compartmental interactions upon in *siNC* vs *siRBFox2* cells. **(G)** Comparison of TAD numbers in *siNC* vs *siRBFox2* cells. **(H-I)** Comparison of TAD size and TAD score in *siNC* vs *siRBFox2* cells. **(J-K)** ADA and APA analyses in *siNC* and *siRBFox2* cells. **(L)** Bar plot showing the number of TADs with +RBFox2 boundaries that were stable, merged, split, disappeared or rearranged upon *RBFox2* depletion.

**Supplementary Figure S9.**
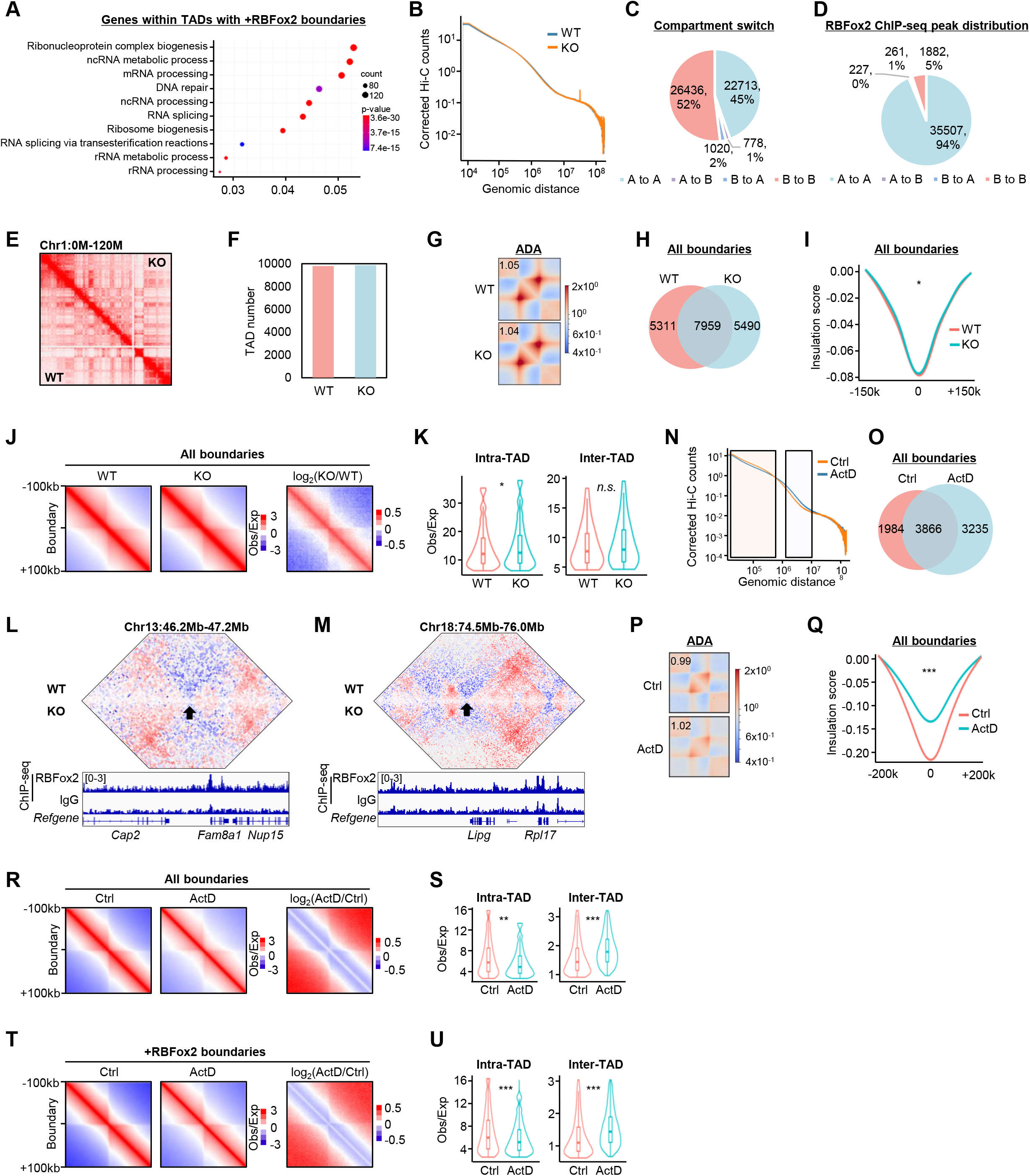
RBFox2 regulation of 3D genome is relevant in mouse myoblast differentiation. **(A)** GO analysis of genes residing in TADs with +RBFox2 boundaries in MBs. **(B)** Contact frequency as a function of genomic distance along the whole genome in WT and KO MBs. **(C)** Pie chart showing the fraction of genome with the designated compartment switching upon RBFox2 knock-out. **(D)** Pie chart showing the percentage of RBFox2 ChIP-seq peaks located at each type of the switched compartments. **(E)** Representative heatmap showing the called compartment at chr10 in KO (upper triangle) vs WT (lower triangle) MB. **(F)** Number of TADs called from WT and KO MBs. **(G)** ADA analysis in WT and KO MBs. **(H)** Dynamic change of all boundaries upon RBFox2 KO. **(I)** Comparison of insulation scores of all boundaries in WT vs KO MBs. **(J)** Aggregate analysis of interactions around all boundaries in WT vs KO MBs. **(K)** Comparison of inter-and intra-TAD interactions around all boundaries in WT vs KO MBs. **(L-M)** Two representative heatmaps showing the decreased interaction frequency upon RBFox2 knock-out. **(N)** Contact frequency as a function of genomic distance along the whole genome in Ctrl vs ActD treated MBs. **(O)** Dynamic change of all boundaries upon ActD treatment. **(P)** ADA analysis in Ctrl and ActD treated MBs. **(Q)** Comparison of insulation scores of all boundaries in Ctrl and ActD treated MBs. **(R)** Aggregate analysis of interactions around all boundaries in Ctrl and ActD treated MBs. **(S)** Comparison of inter-and intra-TAD interactions around all boundaries in **(R)**. **(T-U)** The above analyses were performed on +RBFox2 boundaries in Ctrl and ActD treated MBs.

**Supplementary Figure 10.**
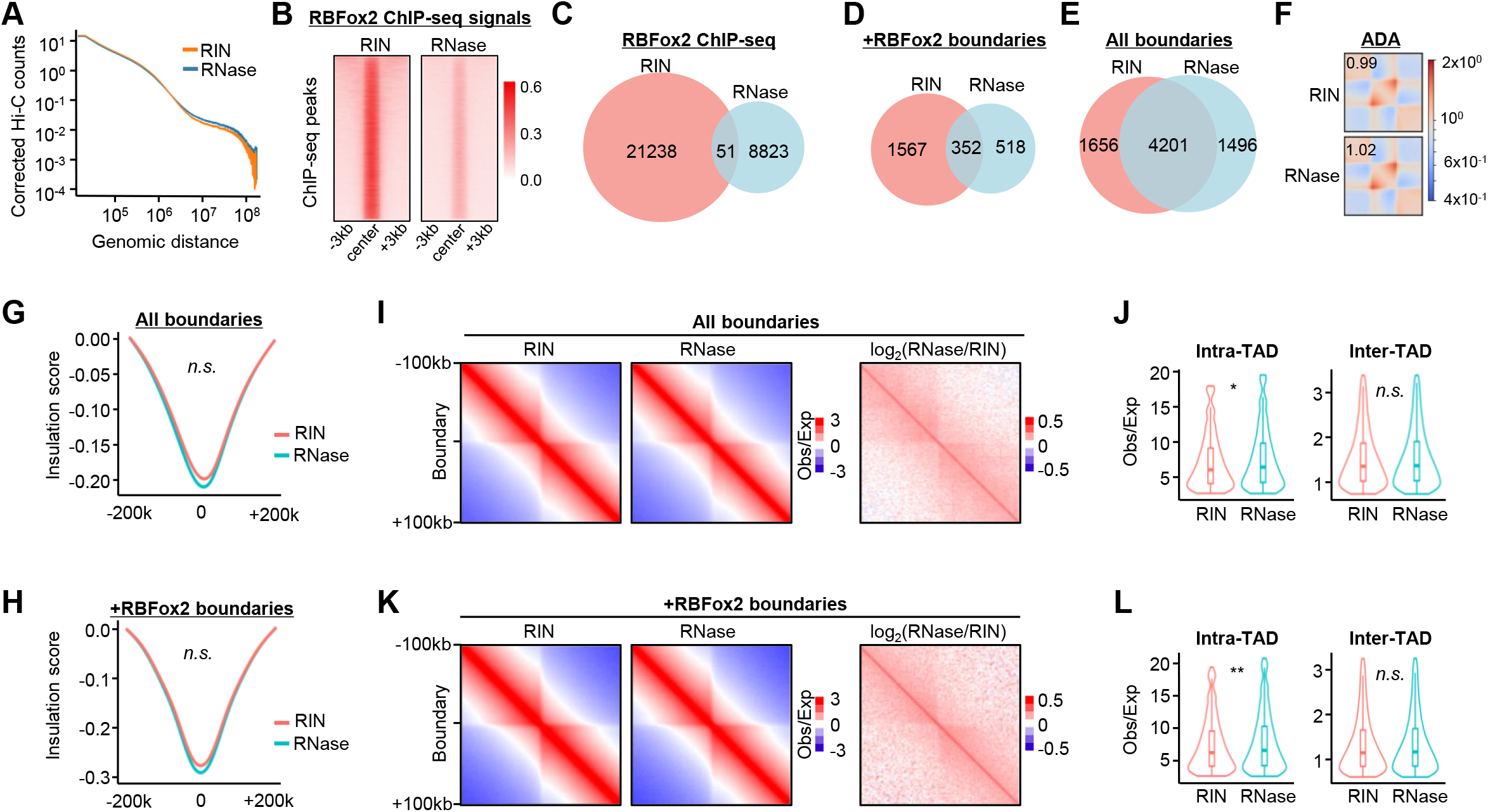
RNase treatment have moderate effect on 3D genome organization. **(A)** Contact frequency as a function of genomic distance along the whole genome in RIN vs RNase treated MBs. **(B)** Heatmap showing the RBFox2 ChIP-seq signal in RIN vs RNase treated MBs. **(C)** Overlapping of RBFox2 ChIP-seq peaks between RIN and RNase treated MBs. **(D)** Dynamic change of +RBFox2 boundaries in RIN vs RNase treated MBs. **(E)** Dynamic change of all boundaries in RIN vs RNase treated MBs. **(F)** ADA analysis in RIN vs RNase treated MBs. **(G)** Comparison of insulation scores of all boundaries in RIN vs RNase treated MBs. **(H)** Comparison of insulation scores of all boundaries in RIN vs RNase treated MBs. **(I)** Aggregate analysis of interactions around all boundaries in RIN vs RNase treated MBs. **(J)** Comparison of inter-and intra-TAD interactions around all boundaries in **(I)**. **(K-L)** The above analyses were performed on +RBFox2 boundaries in RIN vs RNase treated MBs.

